# Rapid and scalable *in vitro* production of single-stranded DNA

**DOI:** 10.1101/558429

**Authors:** Dionis Minev, Richard Guerra, Jocelyn Y. Kishi, Cory Smith, Elisha Krieg, Khaled Said, Amanda Hornick, Hiroshi M. Sasaki, Gabriel Filsinger, Brian J. Beliveau, Peng Yin, George M. Church, William M. Shih

**Author notes:** denotes equal contribution (Dionis Minev, Richard Guerra, Jocelyn Y. Kishi, Cory Smith). Current Address: Department of Genome Sciences, University of Washington, Seattle, Washington, USA.

## Abstract

We present a rapid, scalable, user-friendly method for *in vitro* production of high-purity single-stranded DNA (ssDNA) ranging from 89–3315 nucleotides in length. PCR with a forward primer bearing a methanol-responsive polymer generates a tagged amplicon that enables selective precipitation of the modified strand under denaturing conditions. We demonstrate that the recovered ssDNA can be used for CRISPR/Cas9 homology-directed repair in human cells, DNA-origami folding, and fluorescent in situ hybridization.

---

DNA is instrumental to myriad applications in biological imaging, (bio)nanotechnology, and synthetic biology. Many of the applications rely heavily on the availability of ssDNA. Depending on the required size, scale, and purity, the production of ssDNA can become prohibitively expensive or onerous. Although chemically synthesized ssDNA has become widely available commercially, such DNA has an upper limit of ~200 nucleotides (nt)^1^ in length and often requires additional processing steps to remove impurities. Methods allowing the production of ssDNA through a double-stranded DNA (dsDNA) template via enzymatic processing^2^, micro-bead sequestration^3^, rolling circle amplification^4^, or asymmetric PCR^5^ have been introduced, but are often limited in either complexity of the protocols, scalability, and/or purity of recovered strands. Recently, Palluk et al. have demonstrated a promising enzymatic approach for de novo synthesis of ssDNA^6^. However, this method has yet to demonstrate production of ssDNA >10 nt at high yields. Similarly, autonomous ssDNA synthesis via primer exchange reaction (PER) is currently limited to lengths up to 60 nt^7^. Phagemid-based *in vivo* production of ssDNA can yield biotech-scale quantities of arbitrary sequences, however the method is less amenable to rapid prototyping due to increased lag time between sequence design and strand production^8^. Thus a need persists for methods that allow for fast, user-friendly, scalable, and low-cost *in vitro* production of ssDNA above 200 nt in length.

We present a method that we call <u>Me</u>thanol-<u>R</u>esponsive <u>P</u>ol<u>y</u>mer <u>PCR</u> (MeRPy-PCR), inspired by our previous work of Krieg et al.^9^ (also see Krieg et al.^10^) We create a set of primers bearing a linear polyacrylamide-co-acrylate tag by co-polymerizing a 5’-acrydite-modified primer with acrylamide and sodium acrylate (Fig. 1a, Supplemental Fig. 1, Supplemental Tables 1, 2, Supplementary Protocol 1). The modified primer can include a deoxyuridine (dU), which can be placed anywhere along the sequence and allows the site-specific creation and subsequent cleavage of an abasic site (AB site). We use the polymer-tagged primer in an otherwise standard PCR reaction, resulting in a polymer-tagged amplicon (Fig. 1b, Supplementary Protocol 2) that subsequently allows the selective precipitation (Supplementary Fig. 2) and recovery of both sense and antisense strands away from each other (Fig. 1c, Supplementary Fig. 3). Substitution of a polymer-tagged primer had no noticeably adverse effects on PCR production in terms of strand yield and purity (Supplementary Fig. 4).

**Figure 1:**
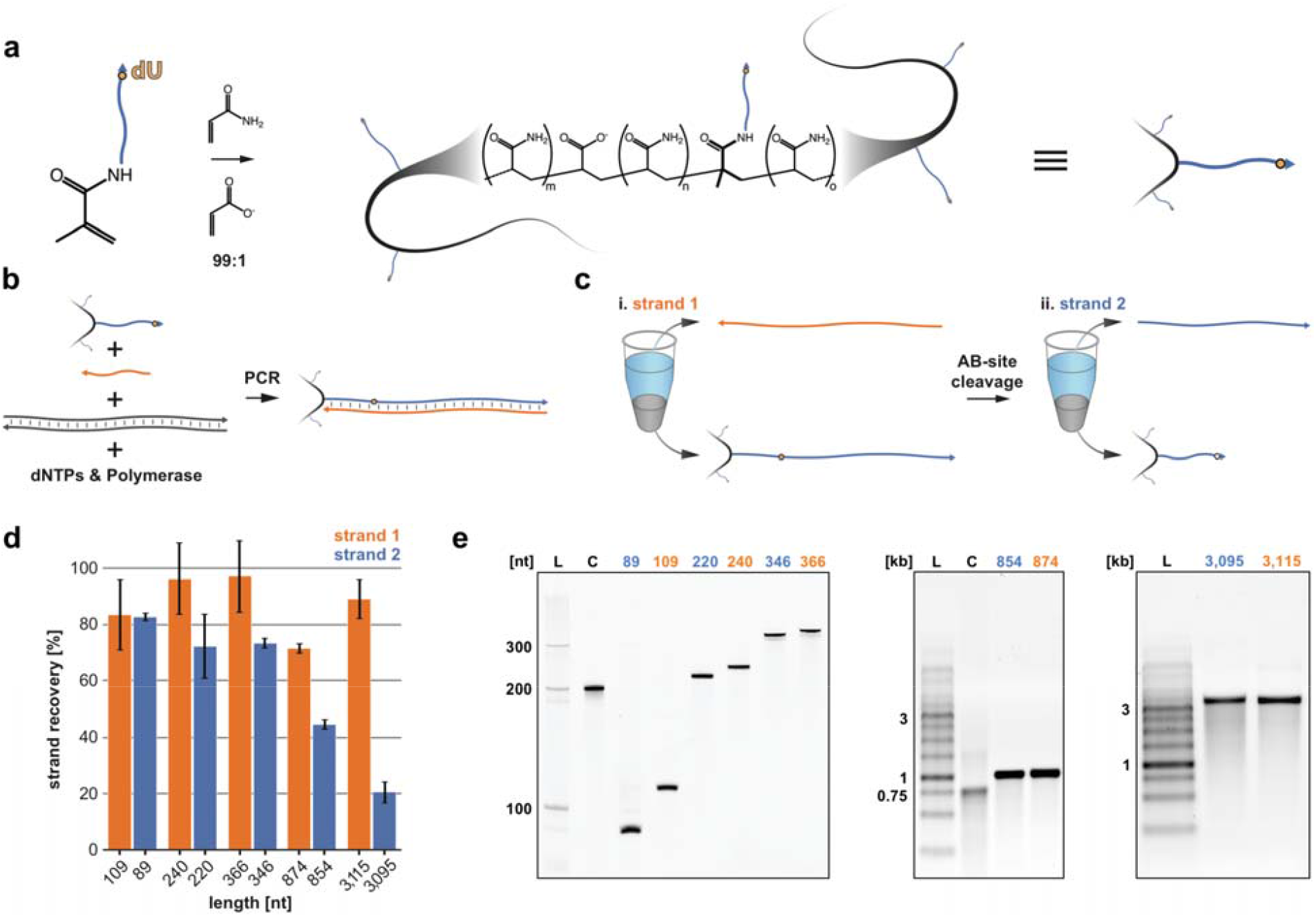
MeRPy-PCR overview and recovery yields for strands 1 (untagged) and 2 (initially tagged) of different amplicon lengths. (a) Production of the polymer-tagged primer. A 5’-acrydite-modified primer is polymerized with acrylamide and sodium acrylate (ratio 99:1) to form a long linear DNA-tagged polymer. (b) MeRPy-PCR procedure following standard PCR guidelines. (c) i. Recovery of strand 1 under alkaline denaturing conditions and methanol precipitation. ii. Recovery of strand 2, after treatment with UDG and DMEDA followed by a methanol precipitation. (d) Recovery yield for strands 1 and 2 of various lengths. Bar graphs denoting the recovery yield (%). Strand recovery yield was determined by the absolute recovered strand output (pmol) relative to MeRPy-PCR input (pmol). Data is shown as mean +/− STD (N=3). (E) Gel electrophoresis of MeRPy-PCR derived ssDNA. Left, denaturing polyacrylamide gel with L – 20 bp Ladder, C – 200mer control from Integrated DNA Technologies (IDT). Middle and right, native agarose gels with L – 1 kb Ladder, C – 750mer control from IDT. MeRPy-PCR derived and commercial ssDNAs were loaded with normalized mass amounts for each gel lane in (E).

After PCR, we first recover untagged strand 1 in a supernatant by performing a denaturing precipitation under alkaline conditions by addition of NaOH to 44 mM final concentration, followed by mixing with one volume equivalent of methanol and then centrifugation at 350–2,000 RCF (Fig. 1ci). We next recover complementary strand 2 by resuspending the precipitated polymer-DNA pellet and incubating it with uracil-DNA glycosylase (UDG) for 15 minutes to excise the dU nucleobase and create an AB site. The AB site is then cleaved by incubating the polymer-DNA solution with 100 mM dimethylethylenediamine (DMEDA)^11^ for 15 minutes, followed by precipitation in 50% methanol to remove the waste polymer-tagged DNA (Fig. 1cii). The procedure is completed within ~45–65 minutes (depending on strand amplicon length), with strand 1 recovery accounting for the first ~15–25 minutes (Supplementary Fig. 3).

We used this method to generate ssDNA ranging from 89–3115 nt in length by amplifying an array of target sequences with MeRPy-PCR and recovering both strands 1 and 2 of each amplicon (Fig. 1d, Supplementary Figs. 5–9, Supplementary Note 1). The strand-recovery protocol was nearly identical for all lengths and templates, apart from slight differences in the alkaline denaturation step for the longest amplicons (see Supplementary Protocol 3). Strand 1 was routinely recovered with a yield of 70% to >90% with respect to the initial MeRPy-PCR amplicon. By contrast, recovery yield of strand 2 was lower as the length of the amplicons increased (see Supplementary Note 2). We recorded absolute yields of ~2.2–12 pmol/100 μL PCR for strand 1 and ~0.5–12 pmol/100 μL PCR for strand 2 (Supplementary Fig. 10, Supplementary Yield Data). It should be noted that the final amount and purity of recovered ssDNA depends on the efficiency and cleanliness of the PCR, therefore PCR optimization may be desirable. Furthermore, we observed that ssDNAs recovered from MeRPy-PCR were of high purity, on par or better than a chemically-synthesized 200mer oligonucleotide after PAGE purification and an enzymatically produced 754mer oligonucleotide purchased from the commercial vendor Integrated DNA Technologies (Fig. 1e).

To demonstrate the utility of MeRPy-PCR generated ssDNA for demand-meeting applications, we show CRISPR/Cas9 mediated homology directed repair (HDR) in human cells, fluorescent in situ hybridization (FISH) imaging, and DNA-origami folding. We picked the untagged strand 1 for each application, based on the higher overall recovery yield and briefer protocol. Each of the three tested applications utilizes ssDNA in varying capacities; DNA origami requires long ssDNA scaffolds (>1 kb)^12–14^, FISH requires a library of >100 nt Cy3-labeled strands to tile specific regions of the genome^15^, and CRISPR/Cas9 directed HDR has seen growing interest in the field to use long ssDNA over dsDNA donors^16–18^, which can be difficult to produce or else prohibitively expensive to purchase at sufficient scale for cell-culture experiments.

For HDR, we assessed the performance of MeRPy-PCR generated ssDNA donors (Supplementary Fig. 11) of varying size, relative to a purchased chemically synthesized 200 nt donor from IDT. The ssDNA donor-mediated HDR removed a stop codon from a broken GFP expression vector, restoring the GFP sequence and expression (Fig. 2a, Supplementary Fig. 12, Supplementary Note 3). We generated 5 different ssDNA donors 200–1000 nt long, only varying the homology-arm length. We produced the ssDNA donors at yields of ~13–34 pmol/100 μL PCR (Supplementary Table 3). The efficiency of HDR was comparable for the different MeRPy-PCR generated ssDNA and the 200 nt chemically synthesized donor.

**Figure 2:**
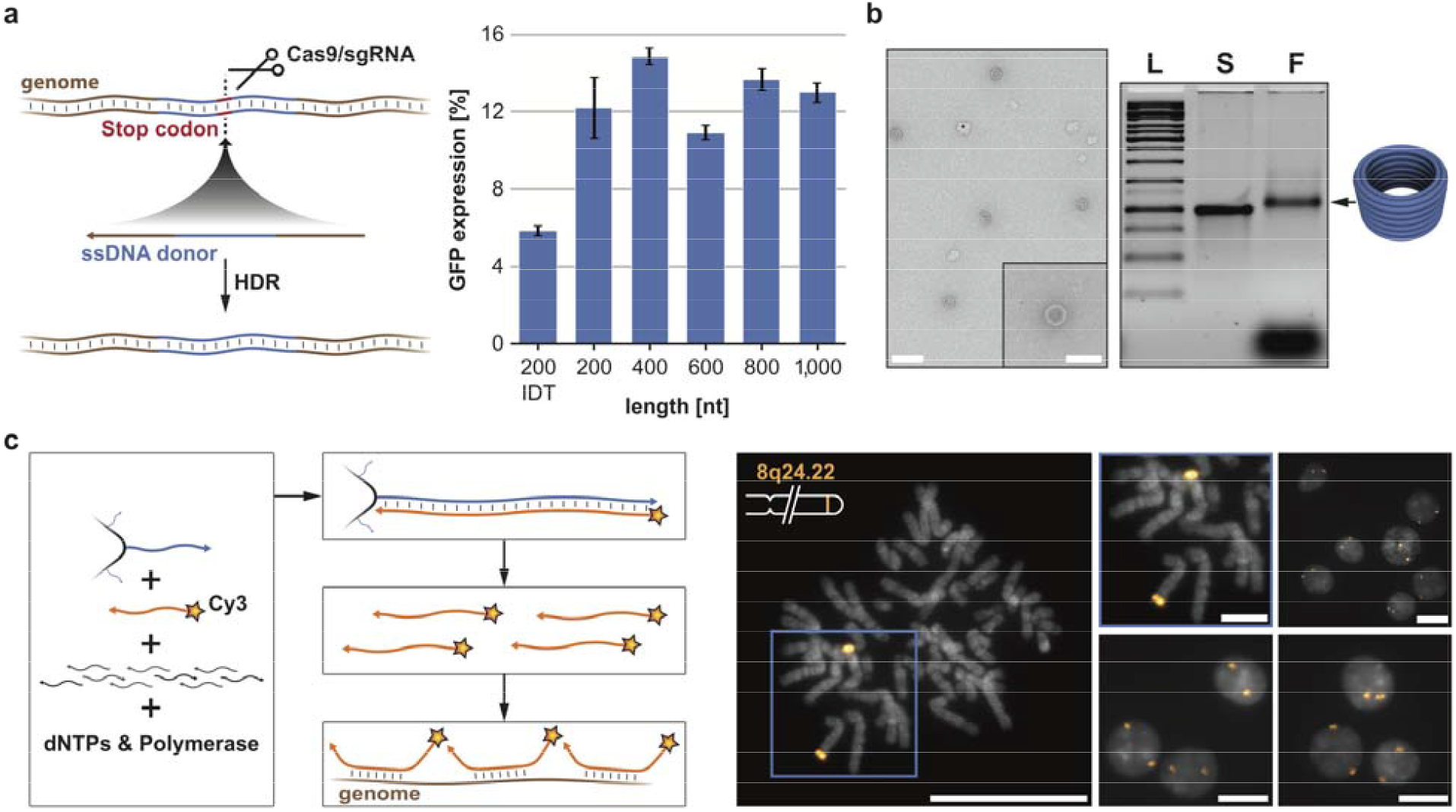
Applications using ssDNA of various lengths. (a) Genome editing in human cells using CRISPR/Cas9. (left) A genomically integrated GFP coding sequence is disrupted by the insertion of a stop codon and a 68-bp genomic fragment from the AAVS1 locus. Restoration of the GFP sequence by HDR with a ssDNA donor sequence results in GFP+ cells that can be quantified by FACS. (right) Bar graph depicting HDR efficiencies induced by MeRPy-PCR derived ssDNAs of different lengths vs. a 200mer chemically synthesized strand from IDT. Data is shown as mean +/− STD (N=3). (b) ssDNA scaffold was generated via MeRPy-PCR from the phage genome, p7308, and used in the folding of a 30 nm DNA origami barrel. Agarose gel electrophoresis (right) shows the purified scaffold strand (S) alongside the folded barrel structure (F). Transmission electron microscopy depicts the folded origami (left). Scale bars denote (left) 100 nm and (right) 50 nm. (c) A library comprising 42,000 probe sequences designed to tile along an 8.4 Mbp region of Human Chromosome 8 was amplified from a small amount of template using MeRPy-PCR with a Cy3-labeled reverse primer and subsequent recovery of fluor-tagged strand 1 library. The generated fluor-labeled ssDNA library was validated in situ on fixed human metaphase spreads and interphase cells. Scale bars denote 20 μm (zoom of metaphase spread scale bar denotes 5 μm).

Next, we tested the ability to produce custom scaffolds for DNA-origami folding. DNA origami is often limited to a defined number of ssDNA scaffolds based on the availability of different M13 phage genomes. There is growing interest in the field for the design and production of new scaffolds that offer a larger range of sequence space^19^. To address this application, we first used MeRPy-PCR to generate a 3315 nt ssDNA derived from p7308 M13 genome (Supplementary Table 4). Using the produced ssDNA scaffold (1.35 pmol/100 μL PCR, Supplementary Table 5) we demonstrated folding of a DNA-origami barrel^20^ (Fig. 2b, Supplementary Figs. 13 and 14).

Finally, we demonstrated the ability to use MeRPy-PCR to generate a large library of FISH probes with a Cy3 modified 5’ end (Fig. 2c, Supplementary Fig. 15). We were able to generate ~70 pmol/100 μL PCR of Cy3-modified ~130 nt FISH probes (Supplementary Table 6), that can successfully be used in imaging a distinct locus of the genome (Chromosome 8q24.22)^14^. As expected by the FISH probe design, we observed two puncta per cell, with the puncta located towards the end of two similarly sized, medium-length chromosomes (Fig. 2c, Supplementary Fig. 16). Use of MeRPy-PCR here highlights the ease with which FISH probes can be generated in sufficient quantities for imaging, obviating the need for expensive and time-consuming purifications.

In summary, we have demonstrated that MeRPy-PCR can be performed without the need for additional optimization beyond that needed for PCR in general, and can be used to recover high yields of both forward and reverse strands, with a briefer protocol and higher yields for the untagged strand. We further demonstrated that the generated ssDNA can be used in a variety of demand-meeting applications in synthetic biology, (bio)nanotechnology, and biological imaging. The short time frame to recover the strands is user-friendly and lowers the bar to rapid in-house production of large quantities of ssDNA. Importantly, the low-cost production of strands via MeRPy-PCR may enable the accelerated exploration of scaffold design space in DNA origami, of genome visualization with FISH, and of the efficiency and off-target effects of single stranded donor DNA in CRISPR/Cas9 HDR.

## Supporting information

Supplementary Sequences

Supplementary Yield Data

## Acknowledgments

The authors would like to thank Christopher M. Wintersinger for fruitful discussions and for providing a 3D rendered figure of the DNA-Origami barrel. This work was funded by support from ONR Award N00014-18-1-2566 and the Wyss Institute at Harvard Core Faculty Award. Research reported in this publication was supported by National Human Genome Research Institute of the National Institutes of Health under award number RM1HG008525, the Uehara Memorial Foundation Postdoctoral Fellowship, a Damon Runyon Cancer Research Foundation Postdoctoral Fellowship (HHMI).

## Author Contributions

Single-stranded DNA production via MeRPy-PCR: D.M., R.G., and W.M.S. designed the experiments. D.M. and R.G. performed the experiments. D.M. and R.G. analyzed the data. DNA origami folding and imaging: D.M., R.G. and W.M.S. designed the experiments. D.M. and R.G. performed the experiments. CRISPR/Cas9 HDR: C.S. designed the experiments. C.S., K.S., and A.H. performed the experiments. C.S., and K.S. analyzed the data. FISH: J.Y.K., H.M.S. and B.J.B. designed the experiments. H.S.M. and B.J.B. designed the sequence library. H.M.S. performed the emulsion PCR. J.Y.K. performed the FISH experiments. D.M., R.G., J.Y.K., C.S. and W.M.S. wrote the paper.

