## Supplementary Information for "Rapid and scalable *in vitro* production of single-stranded DNA"

**Contents**

Materials and Methods	3

Protocols	6

Protocol 1: MeRPy-primer polymerization	6

**Materials**	6

**Part I: Polymerization procedure**	6

**Part IIa: Purification procedure (for tagged primer - TP)**	8

**Part IIb: Purification procedure (for polymer without primer - P)**	10

Protocol 2: General PCR setup	10

Protocol 3: Strand recovery	12

**Materials**	12

**Part I: Strand 1 recovery (native and denaturing precipitations)**	13

**Part II: Strand 2 recovery (UDG/DMEDA strand cleavage)**	14

**Part III: Isopropanol precipitation**	15

Supplementary Figures	16

Supplementary Tables	26

Supplementary Notes	29

References	38

### **Materials and Methods**

**Solvents and reagents for MeRPy-PCR:** All solvents and reagents were purchased from commercial vendors. Methanol (Sigma Aldrich, 322415), Isopropanol (Fisher Scientific, A426P-4), 1,2-Dimethylethylenediamine (Sigma Aldrich, D157805-5G), Molecular biology grade glycogen (Thermo Fisher, R0561), Molecular biology grade acrylamide 40wt% (Sigma Aldrich, 1697-500ML), Sodium acrylate (Sigma Aldrich, 408220-35G), Molecular biology grade tetramethylethylenediamine (Life Technologies Corp, 15524010), Ammonium persulfate (Sigma, A3678-25G). All acrylamide-labeled single-stranded and ultramer/megamer DNA was purchased from Integrated DNA Technologies. All double-stranded DNA templates used were purchased from Twist Bioscience. Taq DNA polymerase (NEB, M0273L), Phusion (NEB, M0531S).

**UV/Vis absorbance:** Nanodrop 2000c Spectrophotometer (Thermo Scientific) was used to record the data.

**Agarose gel electrophoresis (AGE):** UltraPure agarose (Life technologies, 16500500) was used to prepare agarose gels of various percentages. DNA-Origami structures were eluted for 3 hours at 60 V in pre-stained ethidium bromide (Bio Rad, 1610433) gels in 0.5 x TBE buffer containing 11 mM MgCl_2_. Double-stranded and single-stranded DNA was eluted in 0.5 x TBE buffer at 150 V for 1–1.5 hours (depending on the size of the oligo and percentage of gel). Gels were either pre-stained with ethidium bromide (Bio Rad, 1610433) or post-stained with SYBR Gold (Thermo Fisher, S-11494).

**Denaturing and native polyacrylamide gel electrophoresis (dPAGE and nPAGE):** UreaGel System (National Diagnostics, EC-833-2.2LTR) was used to prepare denaturing polyacrylamide gels of various percentages. 40 wt% acrylamide (Fisher Scientific, BP1406-1) was used to prepare native polyacrylamide gels of 20%. Gels were eluted at 200–300 V for 20–40 minutes (depending on the size of the oligo and percentage of gel) and post-stained with SYBR Gold (Thermo Fisher, S-11494) for 20 minutes before imaging. Densitometry analysis of DNA bands was performed with ImageJ (v2.0.0-rc-69/1.52i)^1^.

**Transmission electron microscopy (TEM):** 3 µL of the crude DNA origami folding reaction was applied to a FCF400-CU-50 TEM grid (Fisher Scientific, 5026034) and incubated for 2 minutes, followed by 3 µL of 2% uranyl formate solution containing 25 mM NaOH and air drying. Imaging was performed at 80 kV on a JEOL JEM 1400 plus.

**Polymerase chain reaction (PCR) and DNA origami folding:** A PTC-225 Peltier Thermal Cycler (MJ Research) in conjunction with various thermocycler protocols.

**Metaphase DNA FISH:** The metaphase DNA FISH protocol was developed from ref.’s^2-4^ Human metaphase chromosome spreads (XX 46N or XY 46N, Applied Genetics Laboratories) were denatured in 2×SSC + 0.1% (vol/vol) Tween-20 (SSCT) + 70% (vol/vol) formamide at 70°C for 90 seconds before being immediately transferred to ice-cold 70% (vol/vol) ethanol for 5 minutes. Samples were then immersed in ice-cold 90% (vol/vol) ethanol for 5 minutes and then transferred to ice-cold 100% ethanol for a further 5 minutes. Slides were then air-dried before 25 µL of ISH solution comprising 2×SSCT, 50% (vol/vol) formamide, 10% (wt/vol) dextran sulfate, 40 ng/µL RNase A (EN0531, Thermo Fisher), and MeRPy-PCR generated probe pool at 1.5 μM final concentration was added. Rubber cement was used to seal the hybridization solution underneath a coverslip, and the sample was placed into a humidified chamber inside an air incubator at 45°C overnight. After hybridization, samples were washed in 2×SSCT at 60°C for 15 minutes and then in 2×SSCT at room temperature (2×5 minutes). Samples were then mounted with 12 µL of SlowFade Gold + DAPI (Thermo Fisher S36939) and sealed underneath a coverslip with nail polish before imaging.

**Microscopy:** Imaging of iterative branching samples was conducted on an inverted Zeiss Axio Observer Z1 using a 100x Plan-Apochromat Oil N.A. 1.40 objective. Samples were illuminated by using Colibri light source using a 365 nm or 555 LED. A filter set composed of a 365 nm clean-up filter (Zeiss G 365), a 395-nm long-pass dichroic mirror (Zeiss FT 395), and a 445/50 nm band-pass emission filter (Zeiss BP 445/50) was used to visualize DAPI staining. A filter set composed of a 545/25-nm excitation filter (Zeiss BP 545/25), a 570 nm long-pass dichroic mirror (Zeiss FT 570), and a 605/70 nm band-pass emission filter (Zeiss BP 605/70) was used to visualize Cy3 signal. Images were acquired by a Hamamatsu Orca-Flash 4.0 v3 sCMOS camera with 6.5 µm pixels, resulting in an effective magnified pixel size of 65 nm.

**DNA synthesis and purification:** The 42k library targeting Human Chromosome 8 was ordered from Twist Bioscience and emulsion PCR was performed as previously in ref.’s^2-4^. This emulsion PCR product was diluted to a final concentration of 1.25 pg/µL for subsequent amplification and ssDNA recovery with the MeRPy-PCR protocol containing a Cy3-labeled reverse primer ordered from Integrated DNA Technologies.

**Cas9 directed HDR in human cells with ssDNA:** Human Embryonic Kidney (HEK) 293Ts with a broken GFP expression vector with AAVS1 gRNA targets were obtained from the Church lab that were negative for mycoplasma infection. They were expanded using 10% fetal bovine serum (FBS) in high-glucose DMEM with glutamax passaging at a typical rate of 1:100 and maintained at 37°C with 5% CO_2_. Transfection was conducted using Lipofectamine 2000 (Thermofisher Catalogue # 11668019) using the protocol recommended by the manufacturer with slight modifications outlined below. 24 hours before transfection ~5.0 x 10^4^ cells were seeded per well in a 24 well plate along with 0.5 mL of media. A total of 1 µg of plasmid DNA was transfected using 2 µL of Lipofectamine 2000 per well. The DNA content per well contained 700 ng of hCas9 mixed with 200 ng of gRNA expressing plasmid and 100 ng of ssDNA donor (0.76 pmol for 200bp donor). HDR was measured by percentage of GFP+ through FACS as follows. Three days post-transfection, the cells were harvested using TrypLE and strained before analysis on the BD LSR. Live cell population was gated using SSC and FSC to separate debris and singlets. GFP+ gates were set using a transfected control cell population that did not receive the HDR donor and controls were performed with ssDNA oligo donor transfection alone.

### **Protocols**

#### **Protocol 1: MeRPy-primer polymerization**

##### **Materials**

- Acrylamide-labeled single-stranded DNA (Avoid excessive light exposure)
- Acrylamide (AA) (40 wt% stock solution, stored light protected at 4°C)
- Sodium acrylate (SA) (20 wt% stock solution, stored light protected at 4°C)
- 5x TBE buffer; 500 mM Tris, 500 mM boric acid,10 mM EDTA, pH 8.2
- 10x TE buffer, 50 mM Tris, 1 mM EDTA, pH 8.0
- 5 M NaCl
- Tetramethylethylenediamine (TEMED, stored light protected at 4°C)
- Ammonium persulfate (APS, protected from moisture)
- Nitrogen gas (N_2_)
- Hypodermic needle
- 2 mL glass vial with septum cap
- Cold methanol (MeOH), stored at -20°C
- H_2_O
- 50 mL centrifuge tubes
- 20 mL disposable syringe

##### **Part I: Polymerization procedure**

**Note:** If changing the production scale, ensure that the reaction container (glass vial) has the appropriate size: to achieve efficient N_2_ purging, 1/4 to 2/3 of the container volume should be filled with the reaction solution.

Prepare the following reactions according to the steps listed below:

| **Sample:** | **H2O [µL]:** | **TBE, 5x [µL]:** | **Acrydite tagged primer [µL]:** | **Acrylamide (AA), 40% [µL]:** | **Sodium acrylate (SA), 20% [µL]:** | **TEMED, 5% [µL]:** | **APS, 5wt% [µL]:** | **Total volume [µL]:** |
| --- | --- | --- | --- | --- | --- | --- | --- | --- |
| TP | 285.25 | 100 | 50 | 62.5 | 1.25 | 0.5 | 0.5 | 500 |
| P | 335.25 | 100 | - | 62.5 | 1.25 | 0.5 | 0.5 | 500 |

*Final concentrations in the reaction:*

| **Sample:** | **Final AA concentration [%]:** | **Final SA concentration [%]:** | **Final APS/TEMED concentration [wt%]:** | **Initial concentration of acrydite tagged primer [µM]:** | **Final concentration of acrydite tagged primer [µM]:** |
| --- | --- | --- | --- | --- | --- |
| TP | 5 | 0.05 | 0.005 | 1,000 | 100 |
| P | 5 | 0.05 | 0.005 | - | - |

**TP** – Polymer tagged with primer ssDNA (MeRPy-primer)

**P** – Polymer only

**Steps**

1. Mix first H_2_O, TBE, acrydite tagged primer, AA and SA, then bubble solution 20 minutes with N_2_. **Note:** Pierce the septum cap with the hypodermic needle and make sure the cap is slightly open for bubbling.
I. In the meanwhile, prepare fresh 5 wt% stock solutions of TEMED and APS:
   A. 5% TEMED: 6.5 μL TEMED + 93.5 μL H_2_O
   B. 5 wt% APS: 25 mg APS + 0.5 mL with H_2_O
2. Add TEMED to reaction, swirl vial, then add APS while continuing N_2_ bubbling for 15 minutes.
3. Close septum cap tightly (stops the bubbling) and leave sample attached to nitrogen overnight.
   I. Sample should be highly viscous the next day

- **Pause point:** Samples can be stored at 4°C and purification can be continued at a later date.

##### **Part IIa: Purification procedure (for tagged primer - TP)**

1. Mix nitrogen bubbled solution with 4.5 mL 1X TE
   I. Add 350 µL 1X TE to the vial. Vortex for 1–2 minutes.
   II. Fill a 20 mL disposable syringe with air, attach a needle to the syringe and bend the needle to a ~90° angle, using the needle cover.
   III. Turn vial above 50 mL centrifuge tube upside down. Then use the syringe to push in air to the bottom of the vial, so that the viscous solution is pushed out and drips into the centrifuge tube.
   IV. Add 0.5 mL 1X TE buffer to the emptied vial, vortex for 30 seconds, then repeat step III.
   V. Repeat step IV. 3 times.
   VI. Add another 2.15 mL 1X TE buffer to the centrifuge tube to get to a total volume of 4.5 mL added 1X TE buffer.
2. Vortex or shake at high speed for 10 minutes.
3. Add 25 µL 5 M NaCl and vortex for a few seconds
4. Take out 50 µL aliquot for analysis later (**UP**)
5. Use a syringe: Fill up with 5 mL cold MeOH. **Note:** remove the centrifuge cap and vortex on low speed, in order to avoid spillage.
   I. Slowly add 5 mL, while vortexing. The last 1–2 mL of MeOH should turn the clear liquid turbid, forming a precipitate.
6. Let the sample incubate for 2 minutes on ice.
7. Centrifuge at 150 g at 4°C for 5 minutes and decant supernatant.
   I. Keep 100 µL of supernatant for later analysis (**SN1**)
8. Re-precipitate:
I. Re-suspend pellet by adding 4.5 mL with H_2_O, vortex for 10 minutes, then add 0.5 mL 10X TE buffer and 30 µL 5 M NaCl.
II. Use a syringe: Fill up with 5 mL cold MeOH.
    a. Slowly add 5 mL, while vortexing. The last 1–2 mL of MeOH should turn the clear liquid turbid, forming a precipitate.
III. Spin down again at 150 g for 5 minutes and decant supernatant.
IV. Keep 100 µL of supernatant for later analysis (**SN2**)
9. Add 4.5 mL H_2_O to the pellet and vortex for 10 minutes. Make sure the pellet is perfectly dispersed
10. Add 0.5 mL 10X TE buffer and vortex shortly
11. Divide into aliquots, and store in -20°C freezer (**TP**).

##### **Part IIb: Purification procedure (for polymer without primer - P)**

Steps 1. to 8. are identical. **Note:** the unpurified polymer reaction (UP) and the supernatants 1 (SN1) and 2 (SN2) are not needed for analysis, since there is no DNA present in this sample.

9. Add 5 mL H_2_O to the pellet and vortex for 10 minutes. Make sure the pellet is perfectly dispersed
10. Divide into aliquots, and store in -20°C freezer.

#### **Protocol 2: General PCR setup**

Here we report the generalized PCR mixes and thermocycler protocols for standard Taq polymerase as well as Phusion high-fidelity PCR master mix with HF buffer as these two polymerases generated all the ssDNA needed to carry out the experiments in this report. We also successfully performed this strand purification protocol from PCRs of different polymerases (Hot Start Taq DNA Polymerase and Kapa Taq), highlighting the generalizability of this recovery method. The results for other polymerases are not reported here as the ssDNA generated was not a part of any downstream application or figure reported here.

**Standard Taq polymerase PCR setup**

All PCRs using standard Taq polymerase were prepared using this general protocol:

| **Component** | **100 µL reaction** | **Final concentration** |
| --- | --- | --- |
| Nuclease-free H_2_O | Bring volume up to 100 µL |  |
| 10X Standard Taq buffer | 10 µL | 1X |
| 10 mM (each) dNTPs | 2 µL | 200 µM |
| 10 uM untagged primer | 2 µL | 0.2 µM |
| X uM MeRPy-primer | Variable* | 0.2 µM |
| 10 nM DNA template | 1 µL | 0.1 nM |
| Standard Taq DNA polymerase | 0.5 µL | 2.5 units/100 µL PCR |

*Amount of MeRPy-primer to use depends on stock concentration prepared after polymerization, but the final concentration should be equal to that of the untagged primer.

**Standard Taq polymerase thermocycler protocol**

| **Step** | **Temperature (°C)** | **Time (s)** |
| --- | --- | --- |
| Initial denaturing | 95 | 30 |
| Denaturing | 95 | 15 |
| Annealing | Variable* | 15 |
| Extension | 68 | Variable** |
| Cycle | 29 additional cycles |  |
| Final extension | 68 | 300 |
| Hold | 4 | forever |

*The annealing temperature depends on the exact primer pair used. Annealing temperatures were calculated based on NEB’s Tm calculator’s recommended temperature (generally 5°C below the annealing temperature of the primer with the lower Tm).

**The extension temperature for all lower range amplicons (>500 nt) was 15 seconds, however higher range amplicons followed NEB’s recommend extension time of 1 minute per kilobase.

**Phusion PCR setup**

| **Component** | **100 µL reaction** | **Final concentration** |
| --- | --- | --- |
| Nuclease-free H_2_O | Bring volume up to 100 µL |  |
| 10 uM untagged primer | 5 µL | 0.5 µM |
| X uM MeRPy-primer | Variable* | 0.5 µM |
| 10 nM DNA template | 4 µL | 0.4 nM |
| 2X Phusion master mix | 50 µL | 1 X |

*Amount of MeRPy-primer to use depends on stock concentration prepared after polymerization, but the final concentration should be equal to that of the untagged primer.

**Phusion thermocycler protocol**

| **Step** | **Temperature (°C)** | **Time (s)** |
| --- | --- | --- |
| Initial denaturing | 98 | 30 |
| Denaturing | 98 | 10 |
| Annealing | Variable* | 30 |
| Extension | 72 | Variable** |
| Cycle | X 29–34 additional cycles |  |
| Final extension | 72 | 240 |
| Hold | 4 | forever |

*The annealing temperature depends on the exact primer pair used. Annealing temperatures were calculated based on NEB’s Tm calculator’s recommended temperature (generally at the annealing temperature of the primer with the lower Tm).

**The extension temperature followed NEB’s recommend extension time of 15–30 seconds per kb.

#### **Protocol 3: Strand recovery**

##### **Materials**

- 1% linear polyacrylamide (without DNA primers) in H_2_O
- Methanol (ice cold)
- Basic denaturing buffer (BDB; 0.2 M NaOH, 2 mM EDTA)
- Washing solution (1 part 5 mM Tris- 1 mM EDTA (pH 8), 30 mM NaCl and 1 part MeOH)
- Uracil DNA glycosylase (UDG) and accompanying 10X UDG buffer
- 10X 1,2-Dimethylethylenediamine (DMEDA; 1, buffered to pH 8–9 with acetic acid)
- 1 M sodium chloride (NaCl)
- Isopropanol
- 3 M sodium acetate (NaOAc; pH 5.3)
- Molecular biology grade glycogen
- 70–75% ethanol (EtOH)
- 1X Tris-EDTA (TE)
- H_2_O
- 1.5–2 mL eppendorf tubes

##### **Part I: Strand 1 recovery (native and denaturing precipitations)**

1. Add 1/3 volume of 1 wt% polymer without DNA primers to the PCR sample (i.e. if the initial PCR sample is 300 µL then add 100 µL of 1 wt% polymer)
   I. Prior to adding polymer, can save 5–10 µL for quality control (Raw PCR)
2. Native precipitation: add in 1 volume ice cold methanol (MeOH) to sample and vortex precipitate
3. Incubate for at least 1 minutes, the centrifuge at 350 – 2000 g (4°C) for 5 minutes (Figure 2).
4. Decant or gently pipette the supernatant out
   I. Can save for quality control as supernatant 1 (SN1)
5. Resuspend the sample in H_2_O up to initial sample volume
   I. May want to save 10 µL for quality control as PCR input control for PAGE analysis (Clean PCR)
6. Denaturing step: spike in 0.22 volumes of basic denaturing buffer (BDB; 0.2 M NaOH, 2 mM EDTA; final concentration of 44 mM NaOH) to the sample
   I. For shorter length strands (< 500 nt), vortex then incubate for 1 minute
   II. For mid-length strands (500–1,000 nt), vortex then incubate for 5 minutes
   III. For longer length strands (1,000+ nt), vortex then incubate for 10 minutes
7. Denaturing precipitation: add 1 volume of MeOH, vortex then incubate for 1 minute
8. Spin samples at 350–2,000 g (centrifuge a < 4°C) then transfer supernatant to fresh tube (this supernatant contains strand 1)
   I. Spin supernatant again at 20,000 g (centrifuge a < 4°C) to remove any remaining polymer
   II. Save 10 µL of supernatant for quality control (SN2)
   III. Transfer supernatant to new tube and proceed to isopropanol precipitation
9. Wash pellet with 1 volume of washing solution (1 part 1X TRIS-EDTA, 30 mM NaCl and 1 part MeOH)
10. Vortex for 5 seconds then spin down at 350–2,000 g (4°C) for 5 minutes
11. Decant or gently pipette the supernatant out
12. Resuspend the sample in H_2_O back up to initial sample volume
13. Proceed with sample to strand 2 recovery

##### **Part II: Strand 2 recovery (UDG/DMEDA strand cleavage)**

1. Prepare the following uracil DNA glycosylase (UDG) reaction to create abasic sites from deoxyuridine bases:
   I. 1 volume of sample (all of sample from part one of protocol)
   II. 10X UDG buffer
   III. UDG (0.5 µL per 200 µL sample)
   IV. H_2_O as needed to adjust final volume
2. Incubate at 37°C for 15 minutes
3. Cleave abasic site by spiking in appropriate amount of 10X DMEDA (1 M; pH 8–9; buffered in acetic acid)
4. Incubate at 37°C for 15 minutes
5. Add 0.04 volumes of 1 M NaCl to the sample to aid in polymer precipitation following DMEDA treatment
6. Add 1 volume of MeOH to the sample, vortex, and incubate for 1 minute to precipitate
7. Centrifuge sample 20,000 g (centrifuge a < 4°C) to remove remaining polymer
8. Transfer supernatant to fresh tube and proceed to isopropanol precipitation
   I. Can save 10 µL of supernatant for quality control (SN3)

##### **Part III: Isopropanol precipitation**

1. Set up the following isopropanol precipitation for both strand 1 and strand 2:
   I. Add 0.1 volumes 3 M sodium acetate (NaOAc)
   II. Add 0.8 volumes isopropanol (iPrOH)
   III. Optional: add glycogen as a carrier molecule to improve visibility of precipitated ssDNA (5 µL glycogen per 1 mL of sample
2. Incubate at -20°C for at least 2 hours (usually 12 hours)
3. Spin at 20,000g (centrifuge a <4°C) for 45 minutes and decant supernatant
4. Wash with 70–75% EtOH
5. Spin for 15 minutes at 20,000 g (centrifuge a <4°C) and decant supernatant
6. Optional: repeat wash step
7. Air dry pellet at room temperature or 37°C
8. Resuspend in desired volume of 1X TE or any other preferred buffer
9. Nanodrop samples

### **Supplemental Figures**

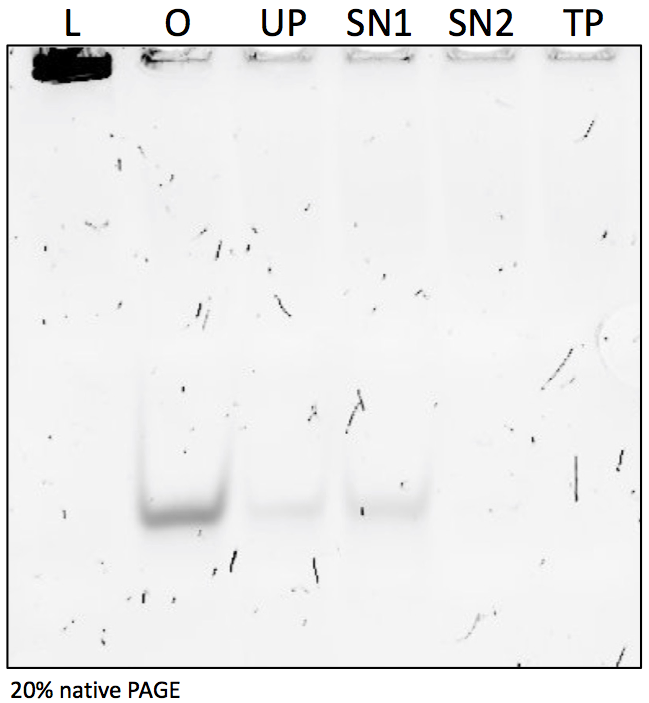

*Supplementary Figure 1: Native polyacrylamide gel electrophoresis of tagged primer and supernatants. (L) 1 kb DNA ladder, (O) acrydite tagged primer oligo, (UP) unpurified polymer tagged primer, (SN1) supernatant 1 after first native precipitation clean up, (SN2) supernatant 2 after second native precipitation clean up, (TP) purified polymer tagged primer. Capture yield was quantified by the amount of primer that was not incorporated into the polymer and thus migrated into the native PAGE.*

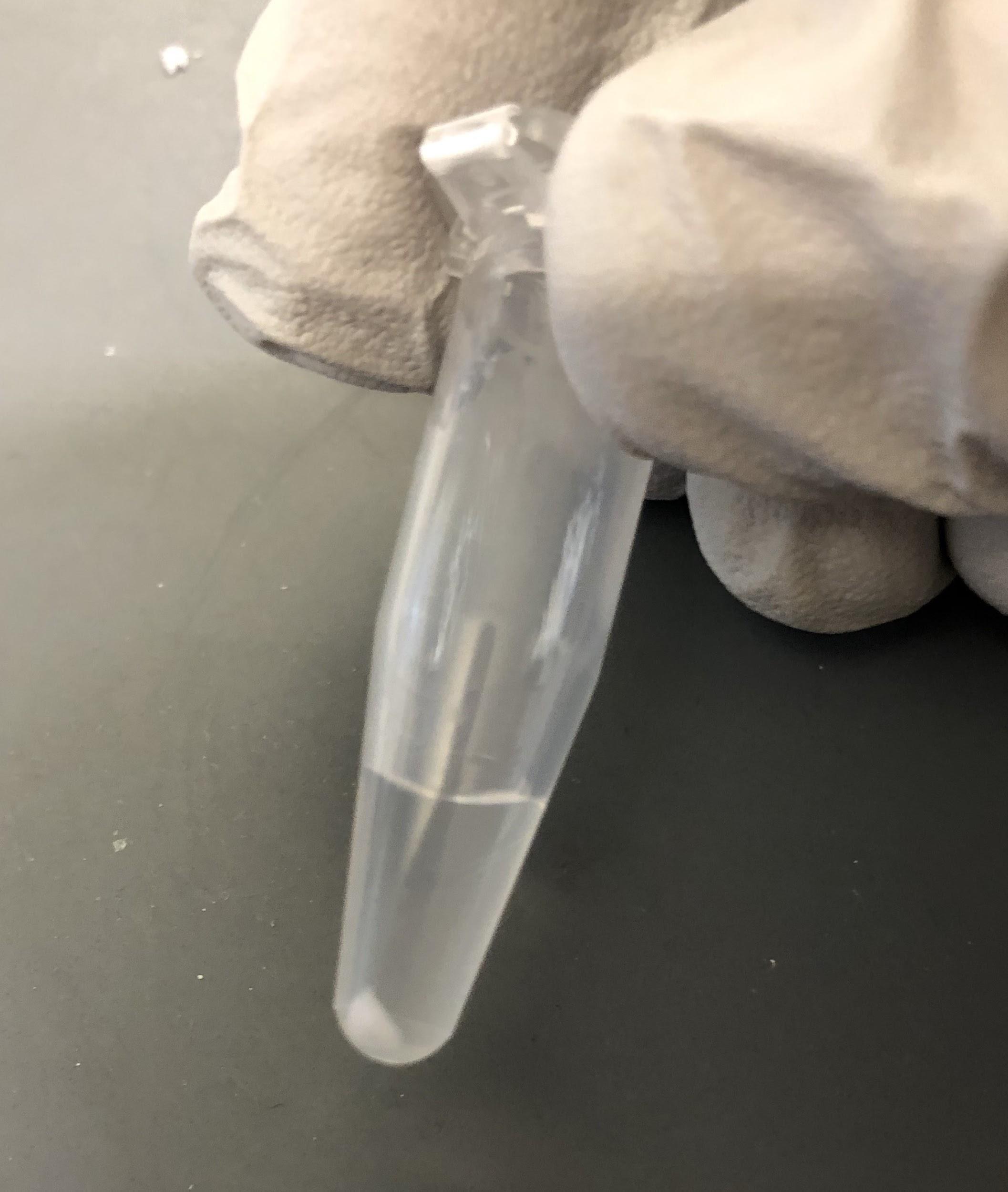

*Supplementary Figure 2: Typical size of a precipitated polymer-pellet with the dsDNA amplicon under native conditions (for 100 µL MeRPy-PCR).*

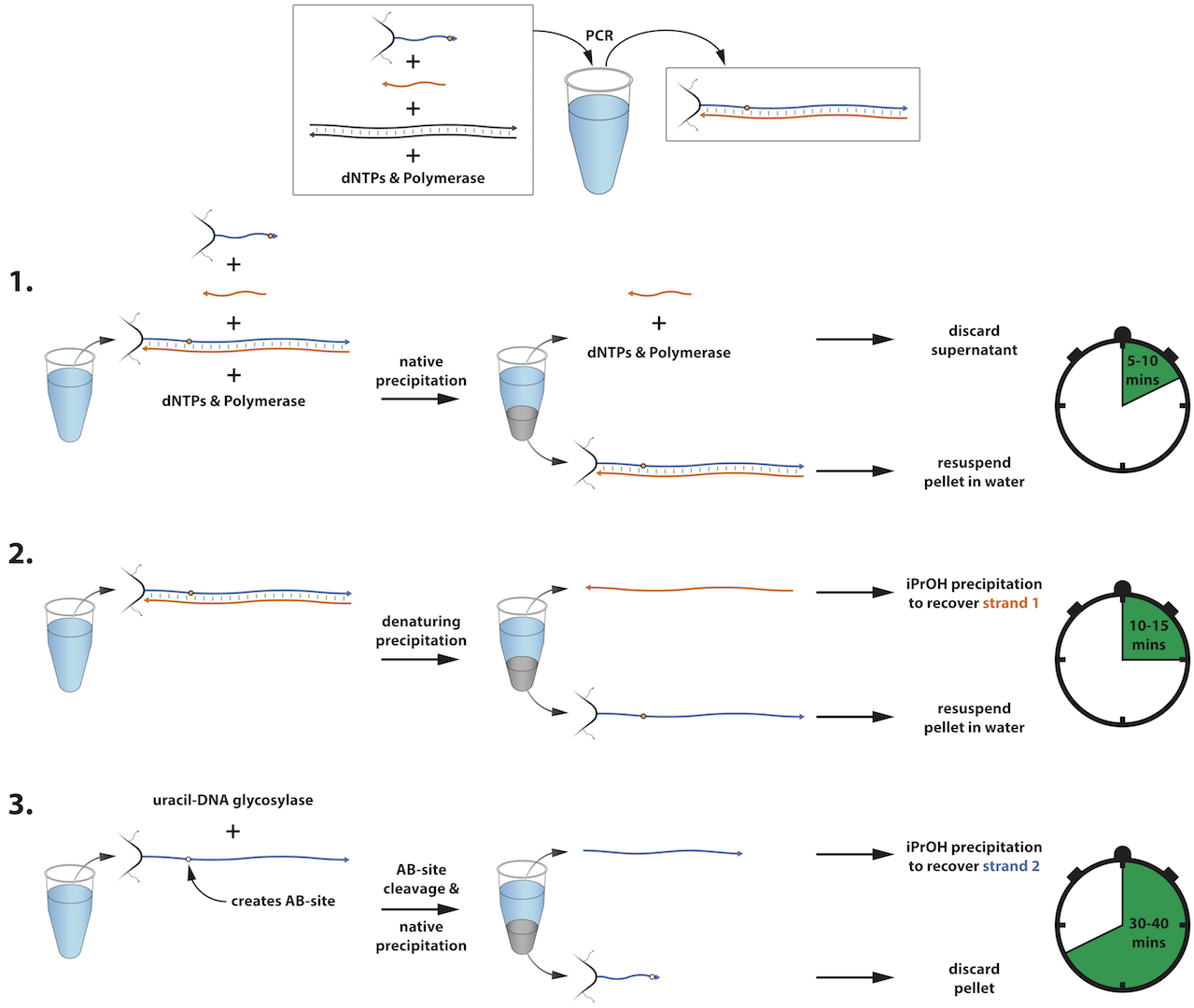

*Supplementary Figure 3: General procedure for the recovery of ssDNA. A standard PCR reaction generates the tagged amplicon. (1.) Addition of linear polymer without DNA handles, water and methanol crashes out the tagged amplicon, followed by discarding the supernatant including reverse primer, dNTPs, and polymerase. (2.) Addition of NaOH and water denatures the amplicon and allows the recovery of strand 1 after the addition of methanol. (3.) Incubation of the tagged strand 2 with UDG, subsequent cleavage with DMEDA and precipitation with methanol allows the recovery of strand 2.*

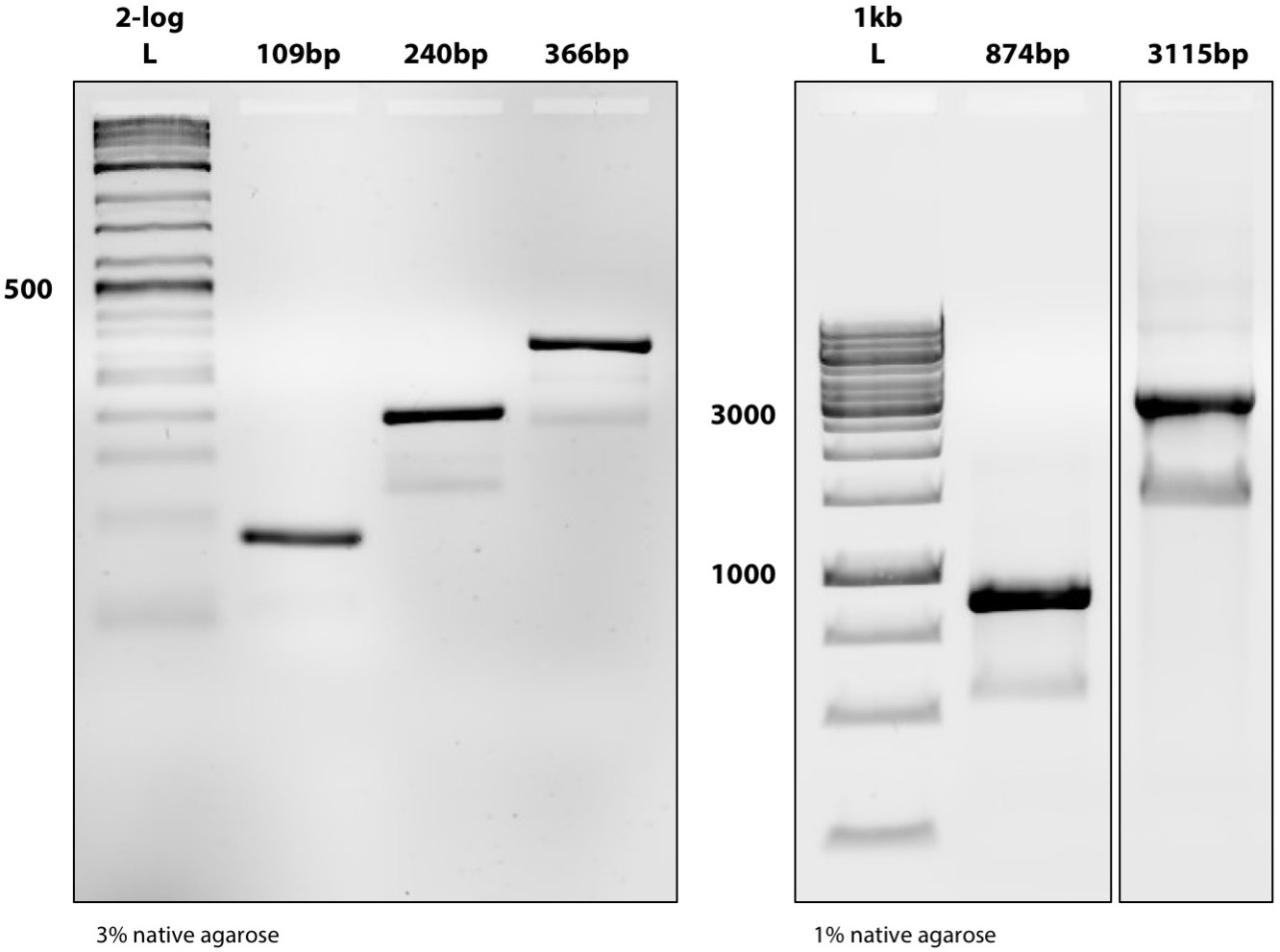

*Supplementary Figure 4: Native agarose gel electrophoresis for standard PCR amplicons (dsDNA) on templates used for MeRPy-PCR ssDNA production (shown in Figure 1, see also Supplementary Yield Data).*

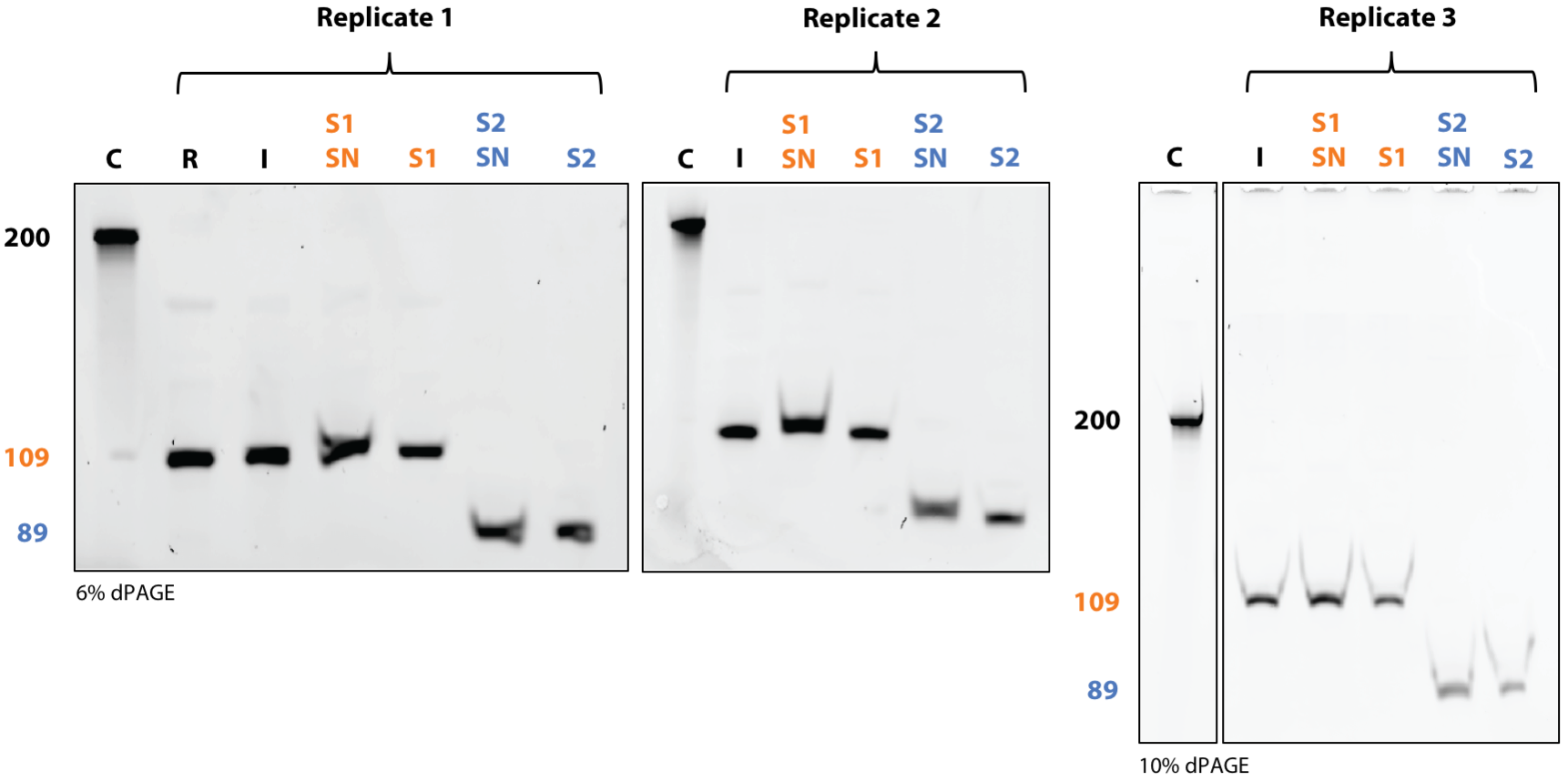

*Supplementary Figure 5: Low range ssDNA recovery with 109 bp template. (C) 200 nt Control oligo from IDT, (R) raw MeRPy-PCR reaction, (I) MeRPy-PCR reaction after first native precipitation, (S1 SN) supernatant after denaturing precipitation of strand 1, (S1) strand 1 after iPrOH precipitation, (S2 SN) supernatant after UDG/DMEDA cleavage of strand 2, (S2) strand 2 after iPrOH precipitation.*

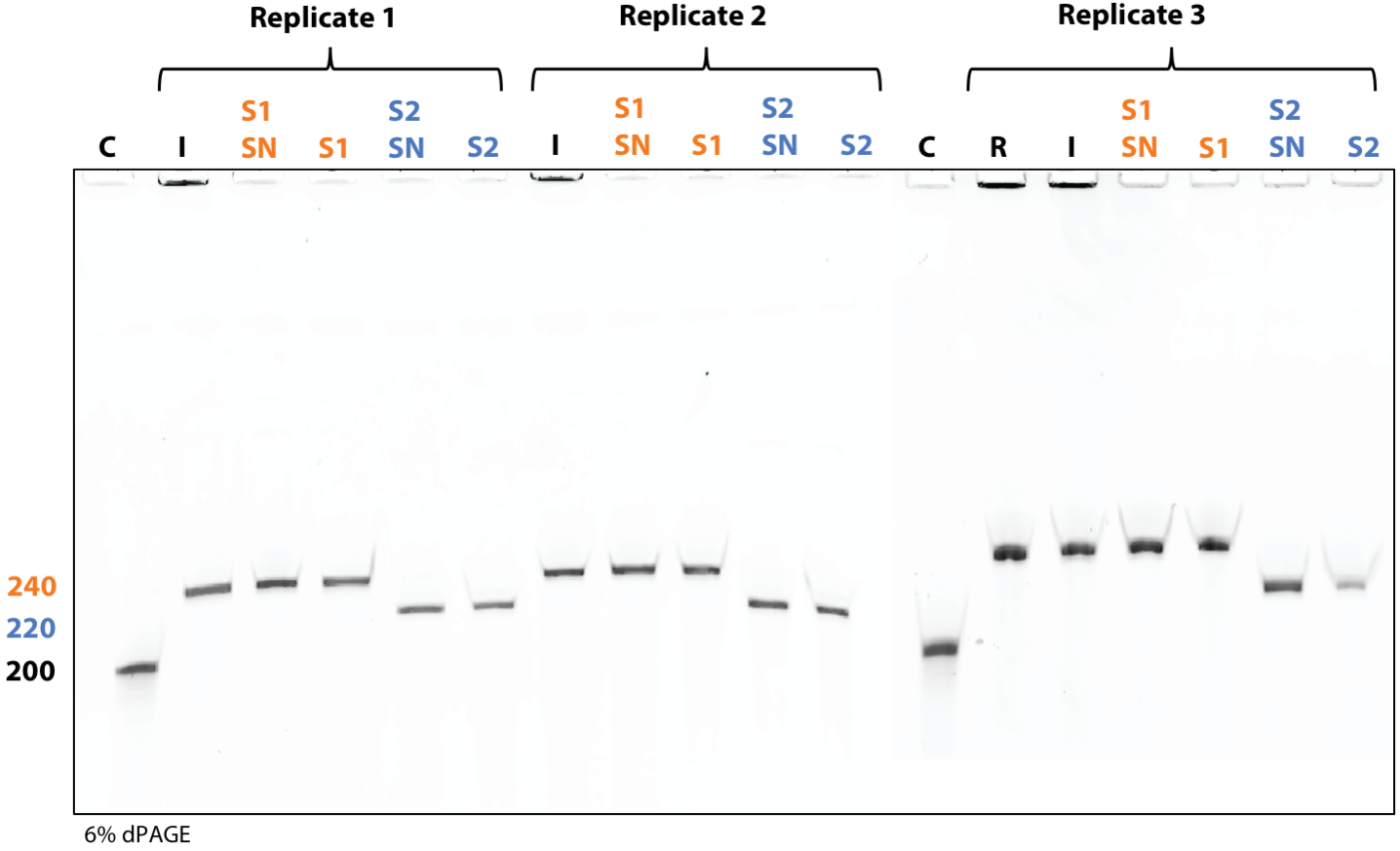

*Supplementary Figure 6: Low range ssDNA recovery with 240 bp template. (C) 200 nt Control oligo from IDT, (R) raw MeRPy-PCR reaction, (I) MeRPy-PCR reaction after first native precipitation, (S1 SN) supernatant after denaturing precipitation of strand 1, (S1) strand 1 after iPrOH precipitation, (S2 SN) supernatant after UDG/DMEDA cleavage of strand 2, (S2) strand 2 after iPrOH precipitation.*

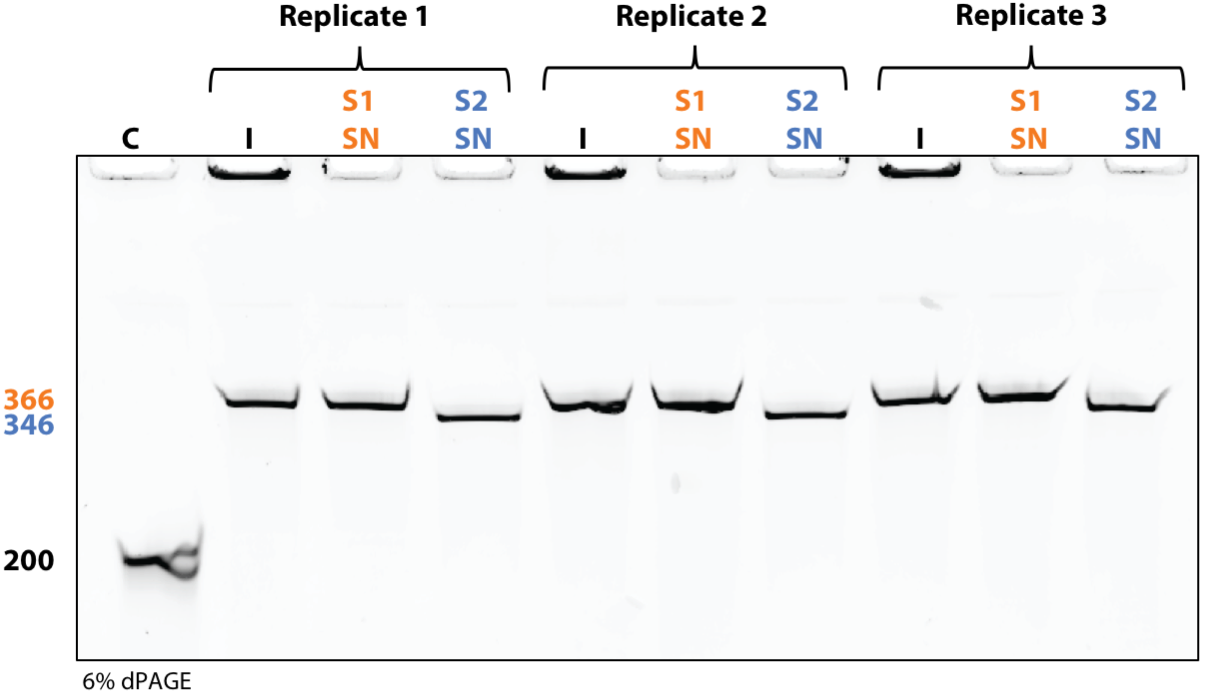

*Supplementary Figure 7: Low range ssDNA recovery with 366 bp template. (C) 200 nt Control oligo from IDT, (I) MeRPy-PCR reaction after first native precipitation, (S1 SN) supernatant after denaturing precipitation of strand 1, (S1) strand 1 after iPrOH precipitation, (S2 SN) supernatant after UDG/DMEDA cleavage of strand 2, (S2) strand 2 after iPrOH precipitation.*

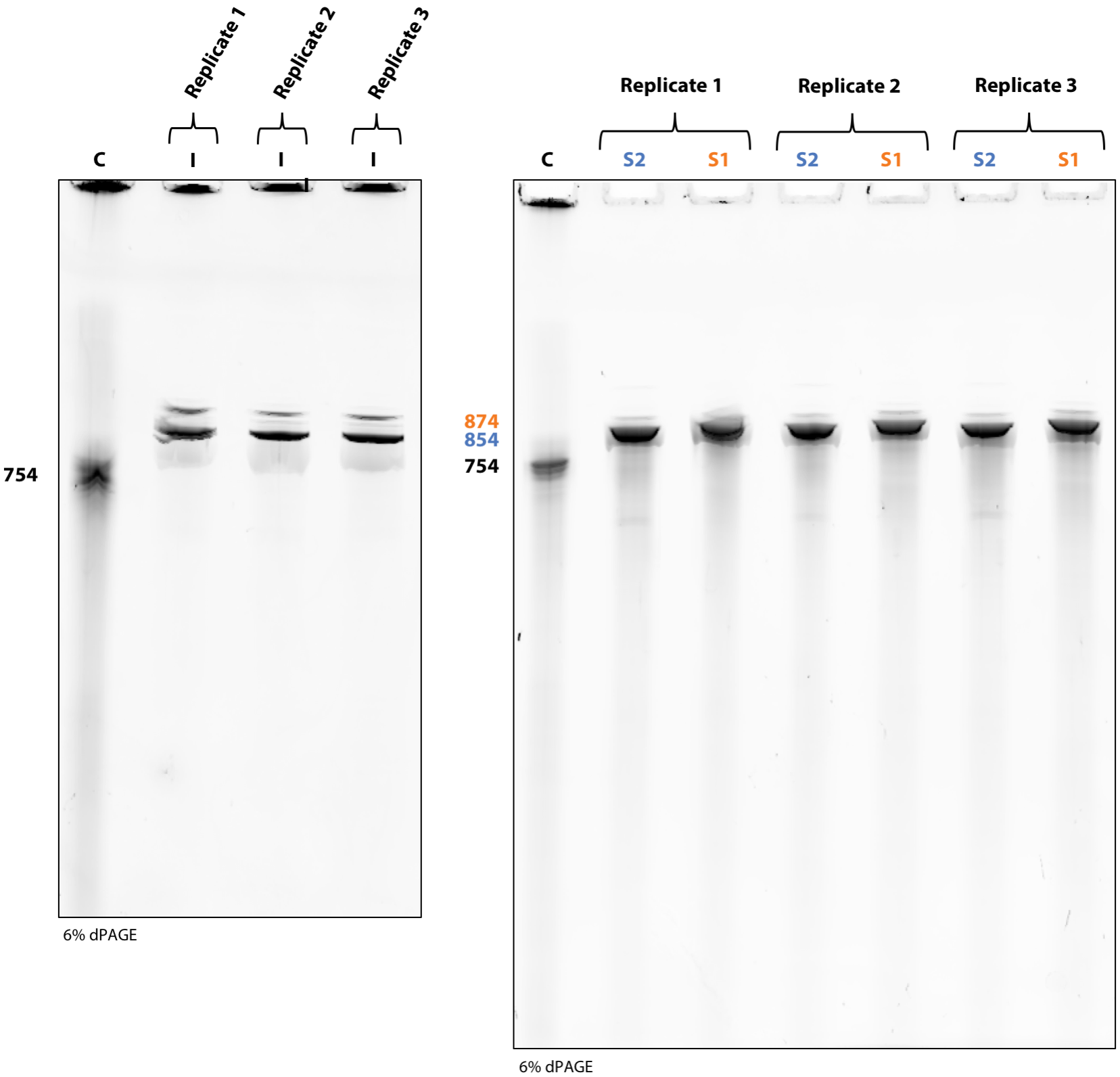

*Supplementary Figure 8: Mid range ssDNA recovery with 874 bp template. (C) 754 nt Control oligo from IDT, (I) MeRPy-PCR reaction after first native precipitation, (S1) strand 1 after iPrOH precipitation, (S2) strand 2 after iPrOH precipitation.*

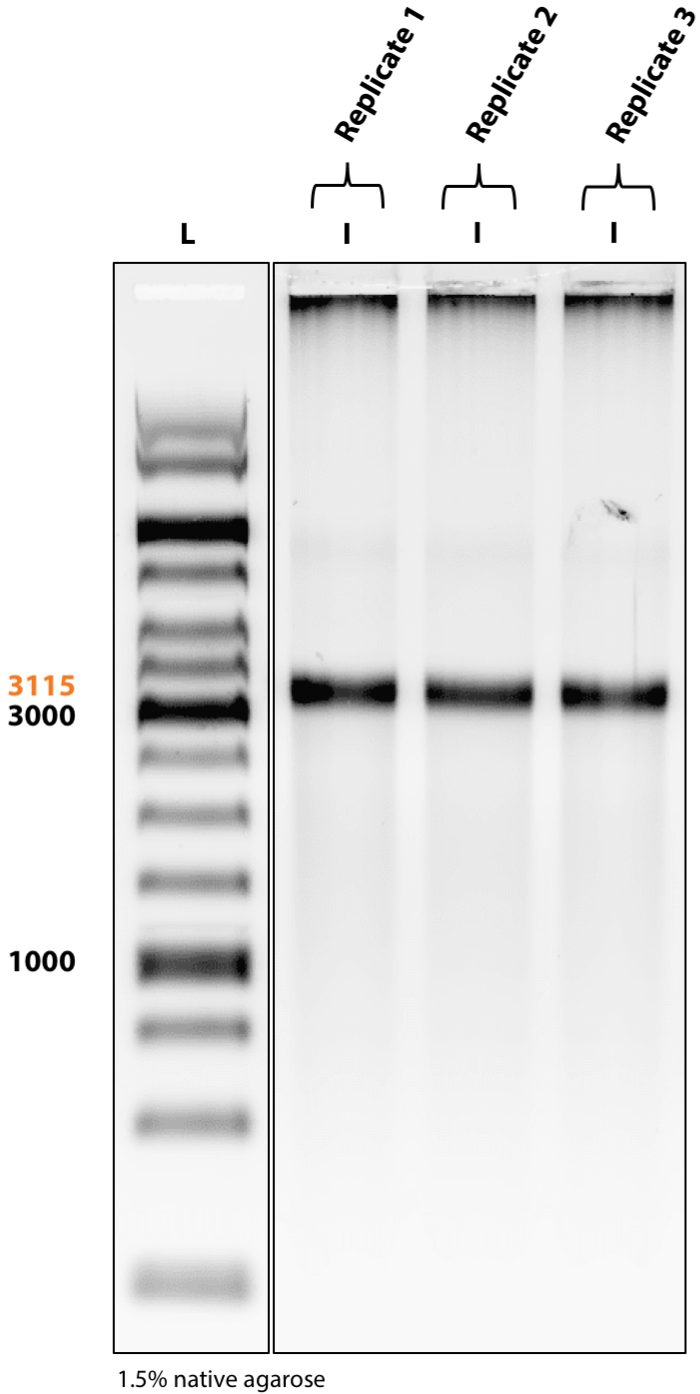

*Supplementary Figure 9: High range ssDNA recovery with 3,115 bp template. (L) 1 kb ladder, (I) MeRPy-PCR reaction after first native precipitation.*

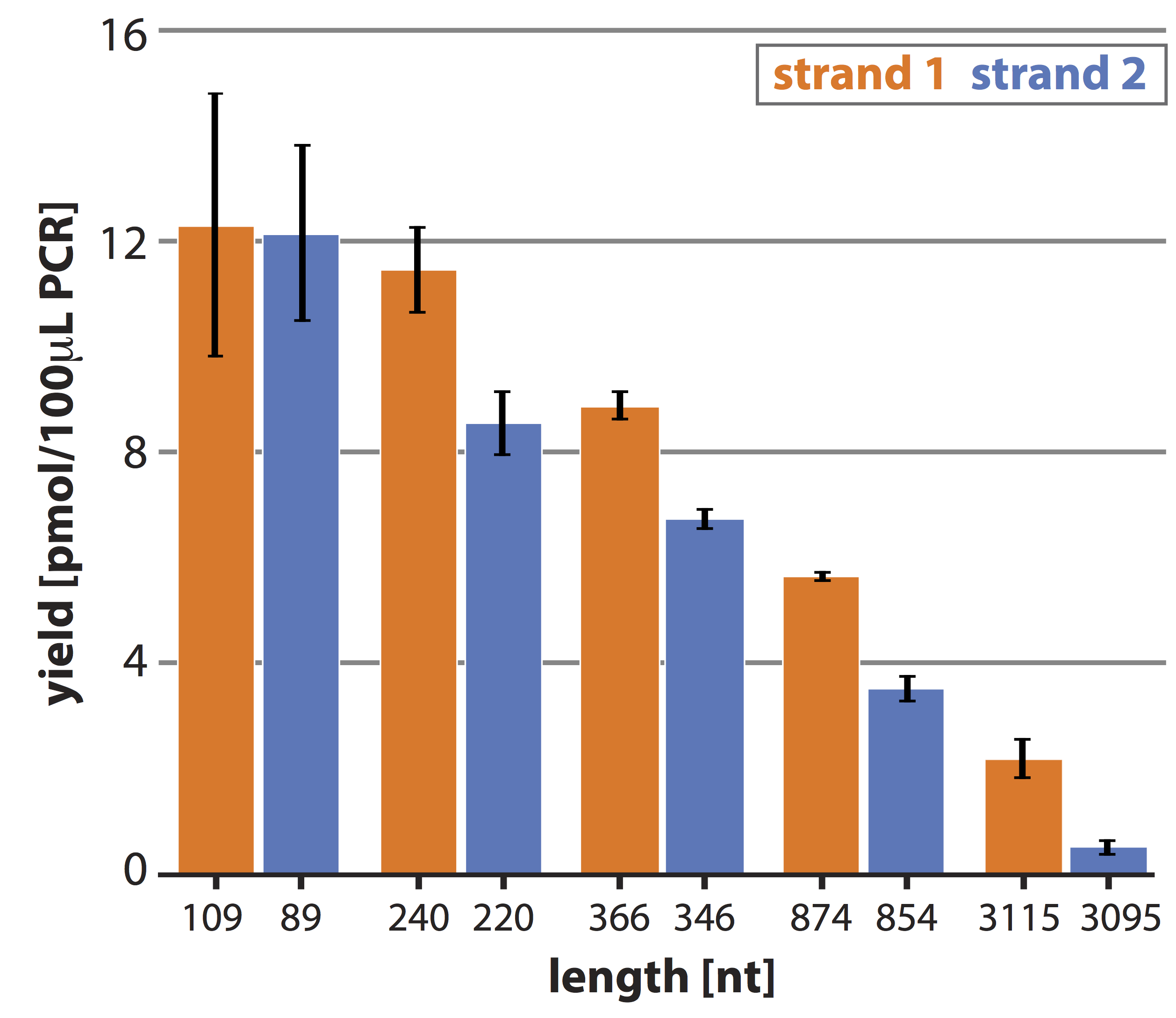

*Supplementary Figure 10: Absolute ecovery yield for strand 1 and 2 of various lengths. Bar graphs denoting the recovery yield (pmol/100 µL MeRPy-PCR reaction). Strand recovery yield was determined by nanodrop after the iPrOH precipitation. Data is shown as mean +/- STD (N=3).*

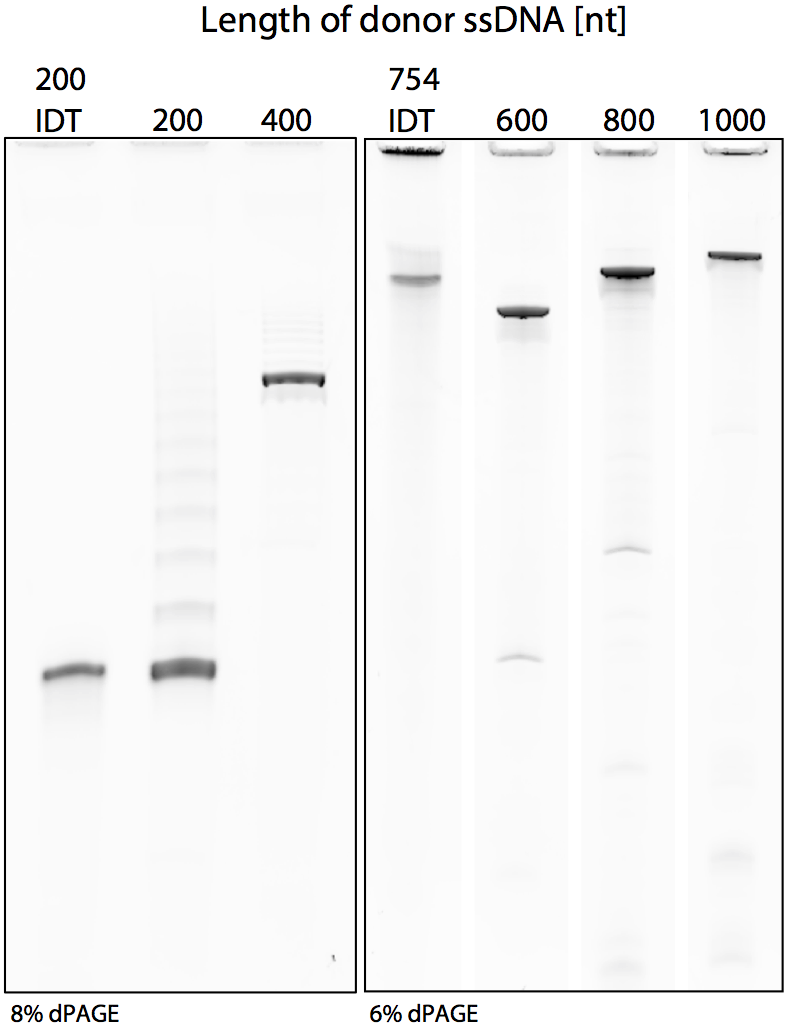

*Supplementary Figure 11: Denaturing polyacrylamide gel electrophoresis of ssDNA donor oligos (made with MeRPy-PCR) used in genome editing in human cells using CRISPR/Cas9. Control lanes on both gels are indicated by 200 and 754 nt from IDT.*

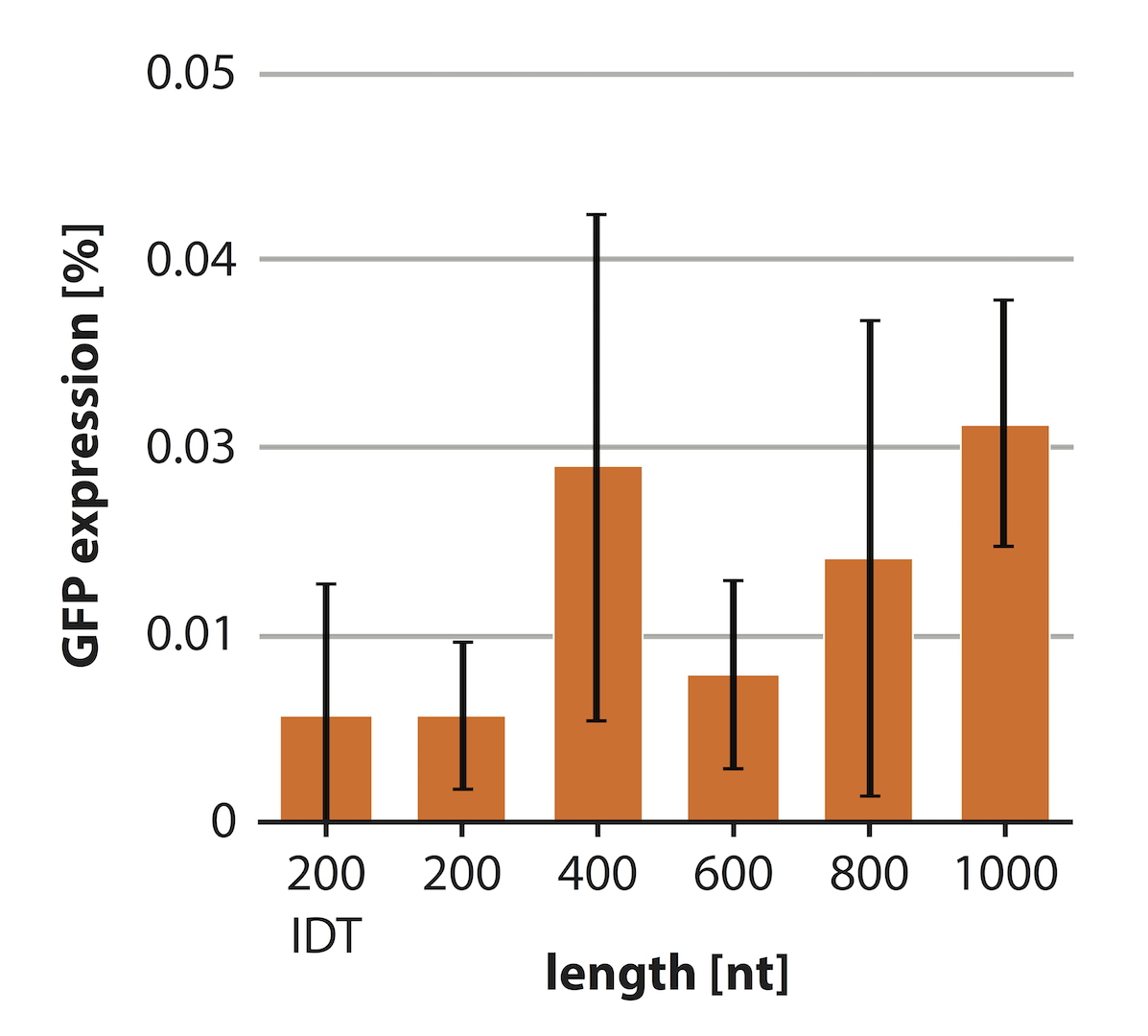

*Supplementary Figure 12: Control experiments for genome editing in human cells using CRISPR/Cas9. Bar graphs showing GFP expression for cell populations transfected with ssDNA oligo donor alone. Data is shown as mean +/- STD (N=3).*

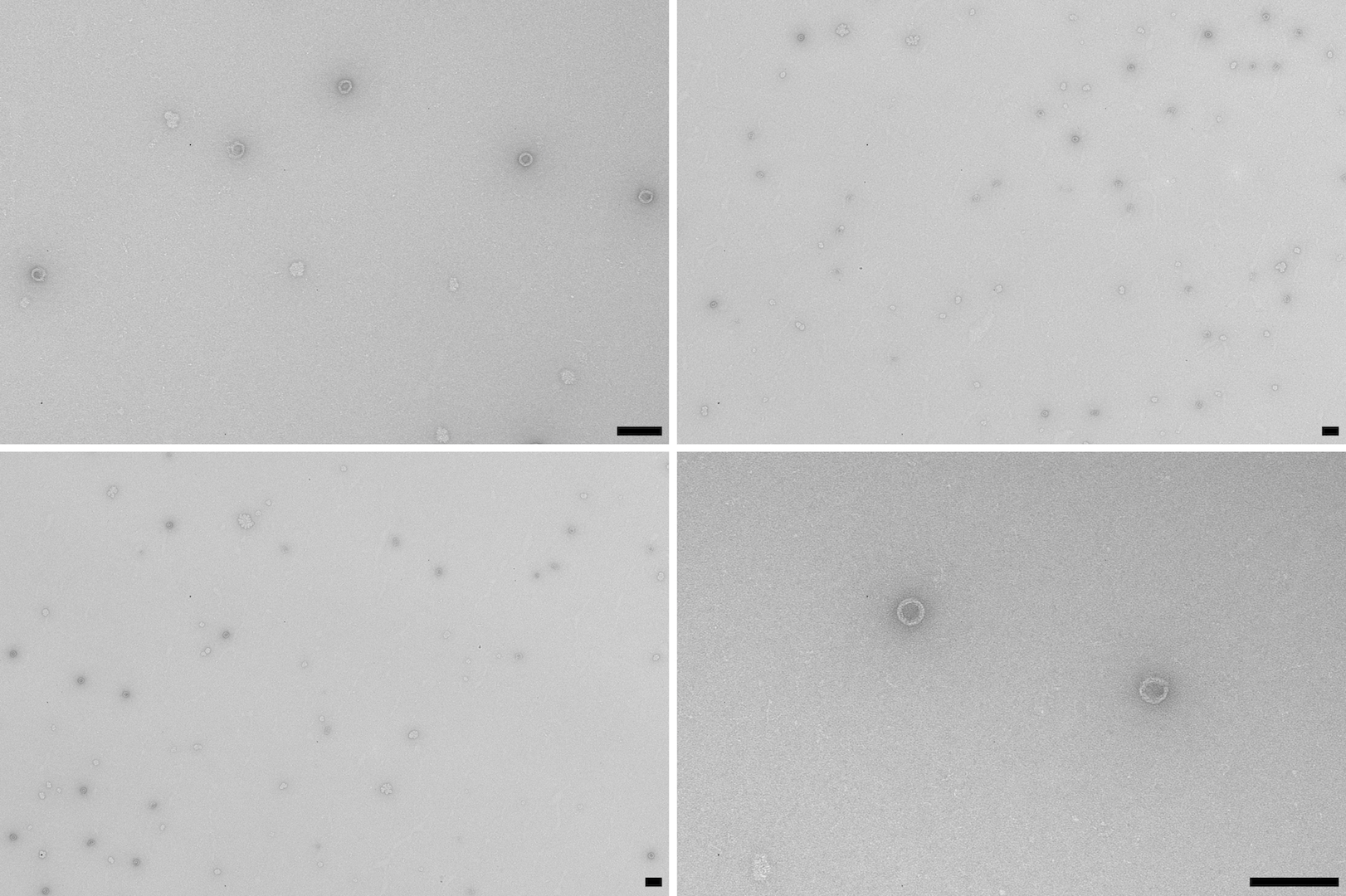

*Supplementary Figure 13: TEM micrographs of DNA origami 30 nm barrel structures. Scale bars denote 100 nm.*

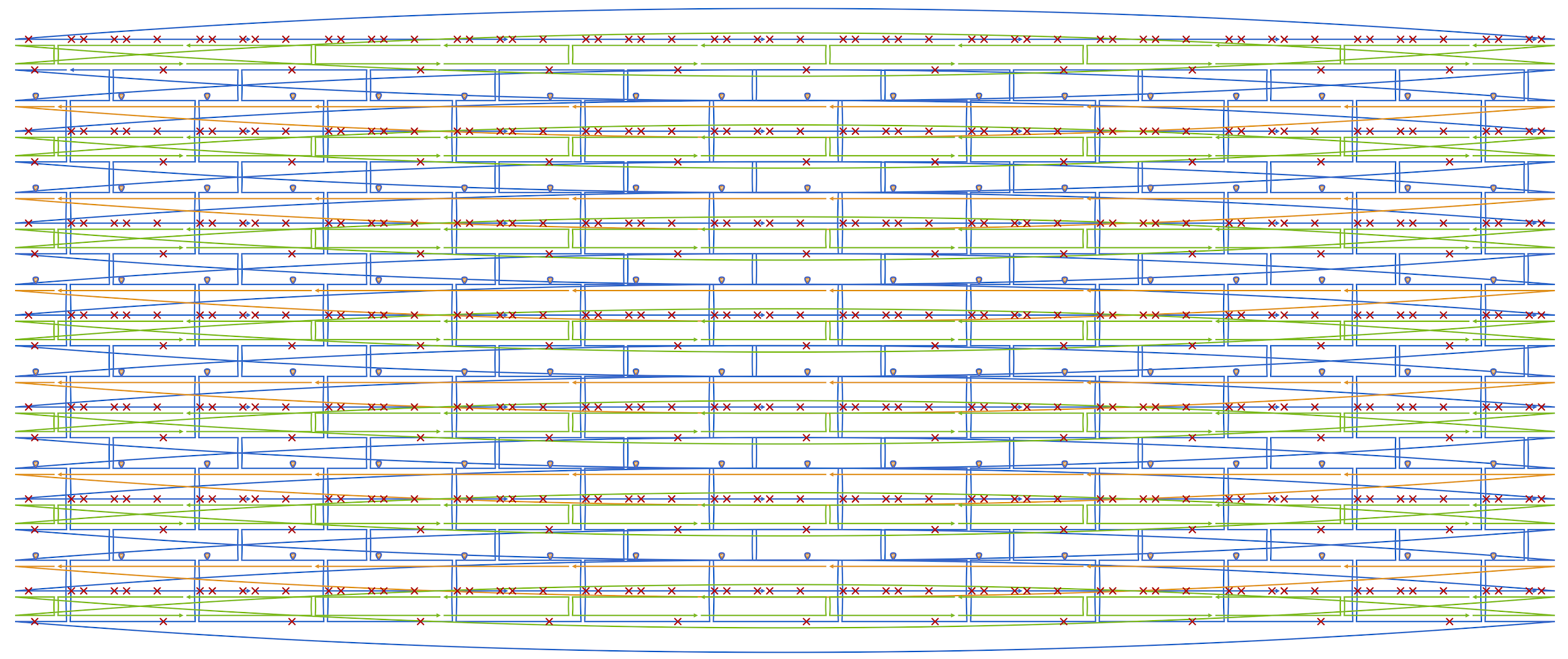

*Supplementary Figure 14: DNA origami strand diagram (generated with caDNAno^5^) of 30 nm barrel.^6^*

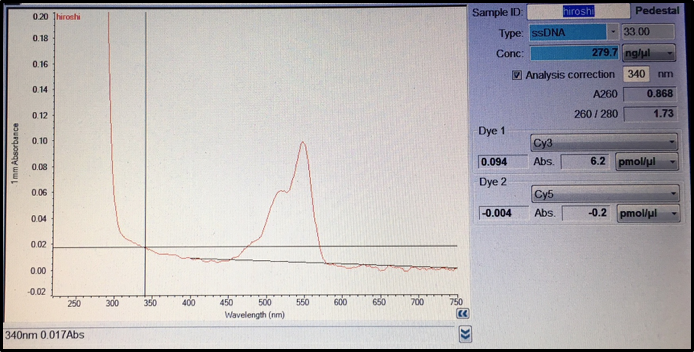

*Supplementary Figure 15: Absorbance spectrum of Cy3-labeled MeRPy-PCR derived ssDNA FISH library.*

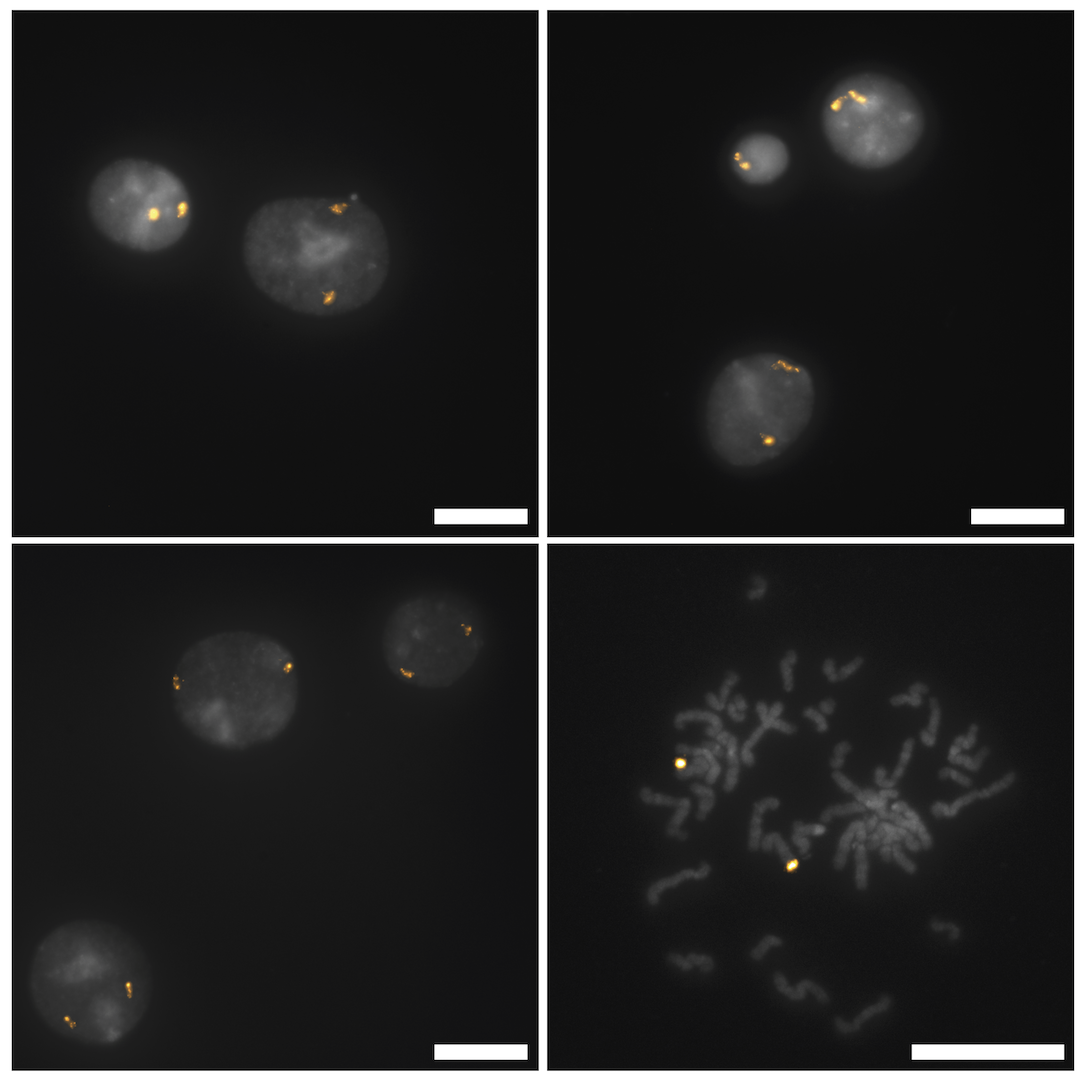

*Supplementary Figure 16:* *A library comprising 42,000 probe sequences designed to tile along an 8.4 Mbp region of Human Chromosome 8 was amplified from a small amount of template using MeRPy-PCR with a Cy3-labeled reverse primer and subsequent recovery of fluor-tagged strand 1 library. The generated fluor-labeled ssDNA library was validated in situ on fixed human metaphase spreads and interphase cells. scale bars denote 20 µm.*

### **Supplemental Tables**

| **Sample:** | **Final concentration (µM):** | **Capture yield (%)** |
| --- | --- | --- |
| TP - polymer tagged primer used in Figure 1 for all templates | 8.823675359 | 88.23675359 |

*Supplementary Table 1: Densitometry analysis results of polymer tagged primer capture yield.*

| **Sample:** | **Nucleic Acid** | **Unit** | **A260 (Abs)** | **Concentration (µM)** | **Capture yield (%)** |
| --- | --- | --- | --- | --- | --- |
| Polymer tagged primer for 200mer | 43.2 | ng/µl | 1.308 | 5.850804486 | 58.50804486 |
| Polymer tagged primer for 400mer | 20.2 | ng/µl | 0.613 | 3.647723784 | 36.47723784 |
| Polymer tagged primer for 600mer | 28.4 | ng/µl | 0.86 | 4.61563465 | 46.1563465 |
| Polymer tagged primer for 800mer | 33.7 | ng/µl | 1.021 | 4.979093716 | 49.79093716 |
| Polymer tagged primer for 1000mer | 23.4 | ng/µl | 0.708 | 3.45729356 | 34.5729356 |
| DNA-Origami scaffold polymer tagged primer | 29.5 | ng/µl | 0.89 | 4.16* | 83.2 |
| FISH probes polymer tagged primer | 53.2 | ng/µl | 1.61 | 8.23 | 82.3 |

*Supplementary Table 2: Capture yield for MeRPy-primers used in Figure 2. Capture yield was determined by nanodrop analysis (preceded blanking with linear polymer of same wt% without DNA handles).*

| **Sample ID** | **Nucleic Acid** | **Unit** | **Strand length (nt)** | **Yield (pmol/100 µL PCR)** |
| --- | --- | --- | --- | --- |
| 200mer | 85.2 | ng/µl | 200 | 34.5 |
| 400mer | 71 | ng/µl | 370 | 15.575 |
| 600mer | 92.8 | ng/µl | 615 | 14.712 |
| 800mer | 137.7 | ng/µl | 805 | 16.782 |
| 1000mer | 110.5 | ng/µl | 975 | 13.425 |

*Supplementary Table 3: Nanodrop results of ssDNA donor oligos used in genome editing in human cells using CRISPR/Cas9.*

| **Ingredient** | **30 nm barrel (µL)** | **Control (µL)** |
| --- | --- | --- |
| 50 mM Tris, 10 mM EDTA, pH 8.0 | 3.4 | 0.96 |
| 60 mM MgCl2 | 4 | 1.13 |
| ss-3315 (50 nM) | 5.5 | 1.6 |
| staple + miniscaf stock (500 nM each) | 8 | - |
| water | 19.1 | 7.61 |
| total volume | 40 | 11.3 |

*Supplementary Table 4: Folding reaction mixture for 30 nm barrel.* *The components were mixed together and then annealed over the course of 20 hours (80°C (10 minutes), 55°C 🡪 45°C (18 hours, 1 hour 48 minutes / °C), 45°C 🡪 25°C (1 hours), 4°C (hold)). Samples were analyzed via agarose gel electrophoresis (0.5 x TBE; 11 mM MgCl_2_; pre-stained with EtBr; resolved at 60 V for 3 hours). Upon confirmation of successful folding, the samples were analyzed by negative staining transmission electron microscopy.*

| **Sample ID** | **Nucleic Acid** | **Unit** | **Strand length (nt)** | **Yield (pmol/100 µL PCR)** |
| --- | --- | --- | --- | --- |
| ssDNA scaffold for DNA origami | 306.1 | ng/µL | 3315 | 1.35 |

*Supplementary Table 5: Nanodrop results of ssDNA scaffold used in DNA origami folding of the 30 nm barrel structure.*

| **Sample ID** | **Nucleic Acid** | **Unit** | **Strand length (nt)** | **Yield (pmol/100 µL PCR)** |
| --- | --- | --- | --- | --- |
| FISH library | 281.6 | ng/µl | ~130 | 70.40966 |

*Supplementary Table 6: Nanodrop results of ssDNA probes used in FISH imaging.*

### **Supplemental Notes**

**Supplementary Note 1:** Recovery yield of strand 1 and 2

Data shown in Figure 1 is calculated based on the triplicate MeRPy-PCR results shown below. The nanodrop results (calculated into pmol) from the recovered strands 1 and 2 after iPrOH precipitation were compared to the densitometry analysis (Supplementary Yield Data) of the MeRPy-PCR reaction after the first native precipitation. Amplicons generated with MeRPy-PCR are tagged and unable to migrate into the gel. In order to obtain MeRPy-PCR yields, the samples had to be denatured and separated on denaturing polyacrylamide gel electrophoresis, allowing strand 1 to migrate into the gel. The 3,115 bp MeRPy-PCR amplicon was denatured with formamide and subsequently eluted on a native agarose gel.

**Supplementary Note 2:** Effect of NaOH denaturation on the recovery yield of strand 2

We have noted that extended exposures of the linear polyacrylamide to the denaturing buffer (NaOH) can cause irreversible polymer damage. Incubation times beyond 10 minutes leads to complete destruction of the polymer and its inability to precipitate from solution with MeOH. We have also observed that longer incubations with denaturing buffer lead to decreased strand 2 recovery, likely caused by the additional damage to the polymer. We hypothesize that this is why we see diminishing strand 2 recovery as the strand length increases; longer strands require longer denaturing incubations and therefore result in lower recovery yields of strand 2. This hypothesis is in accordance with our results in Figure 1 as we see consistent strand 2 yields for the lower range strands as they all follow the same denaturation protocol. However, we see a decrease in yield for our mid-length strand and an even further decrease in yield for our longer length strand.

**Supplementary Note 3:** Fluorescence-activated cell sorting (FACS) scatter plots

Live cell population was gated using SSC and FSC to separate debris and singlets. GFP+ gates were set using a transfected control cell population that did not receive the HDR donor and controls were performed with ssDNA oligo donor transfection alone.

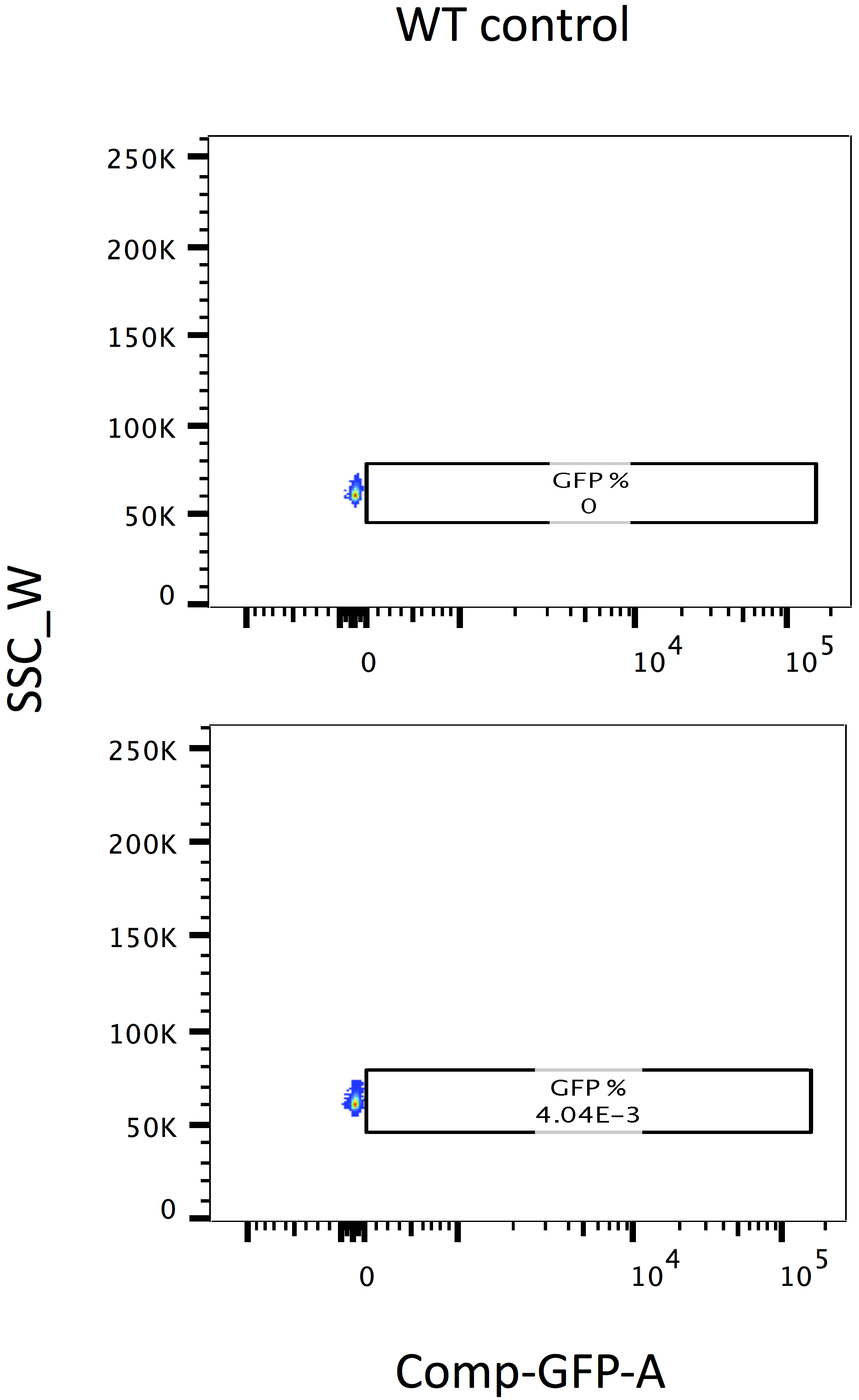

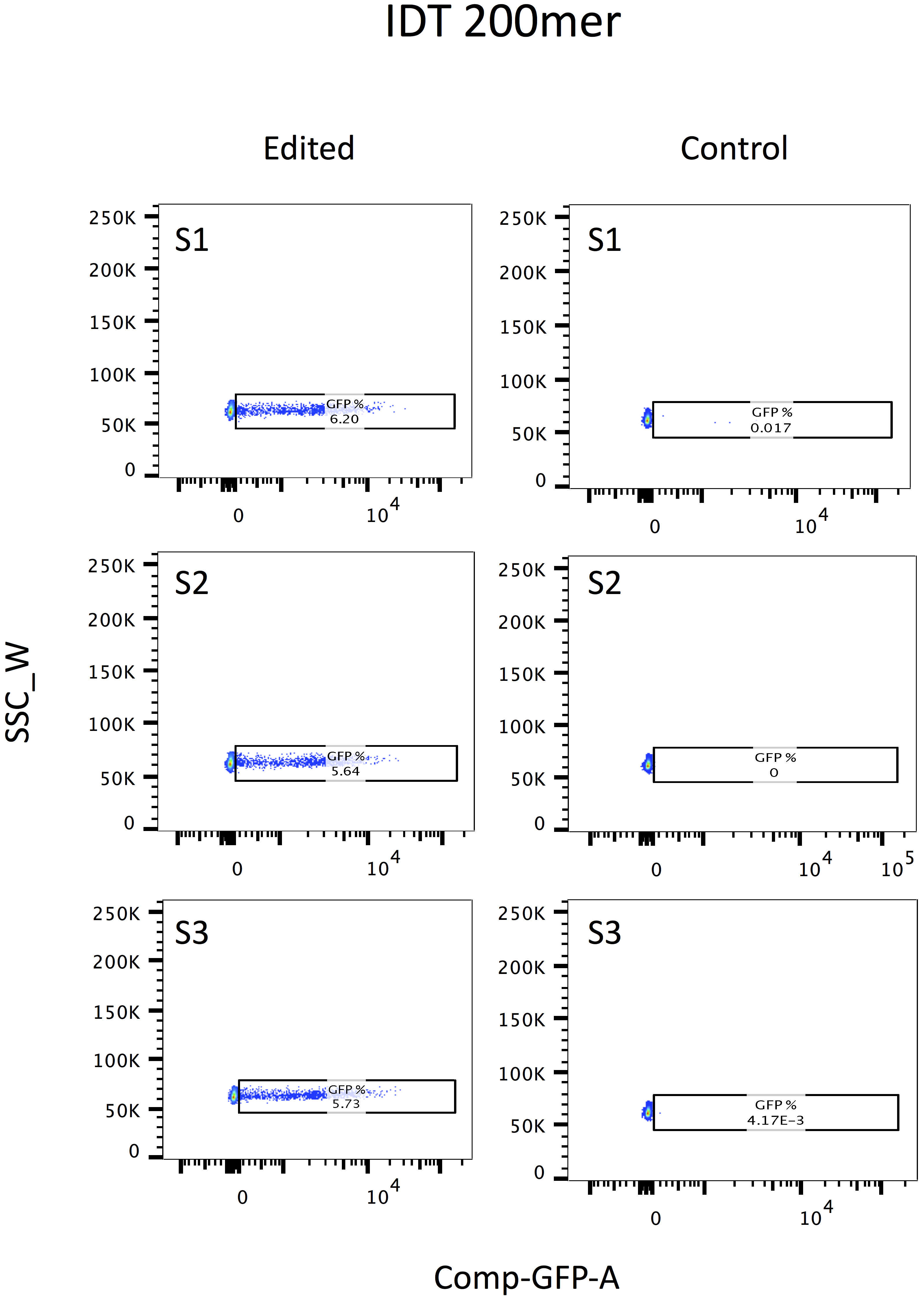

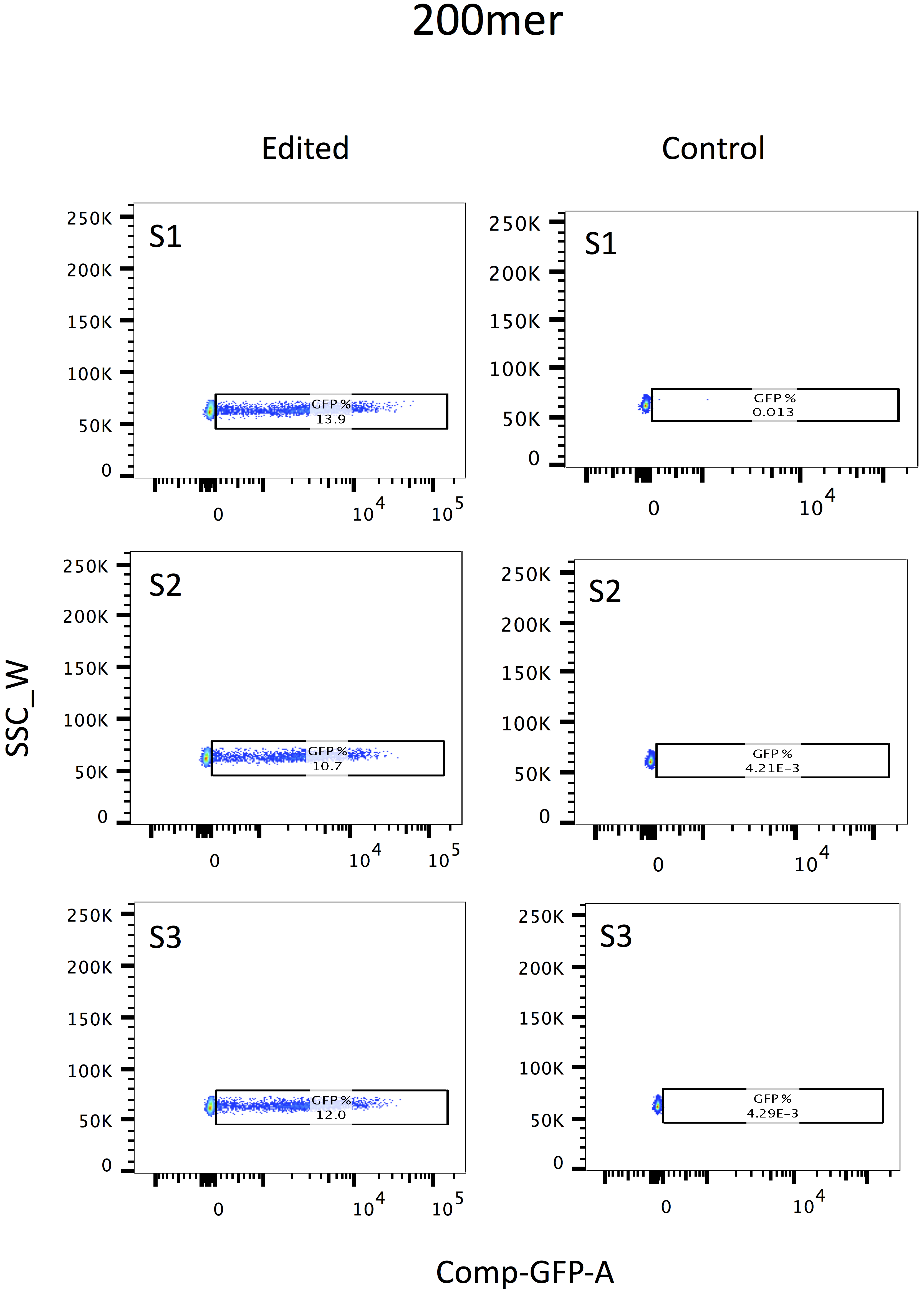

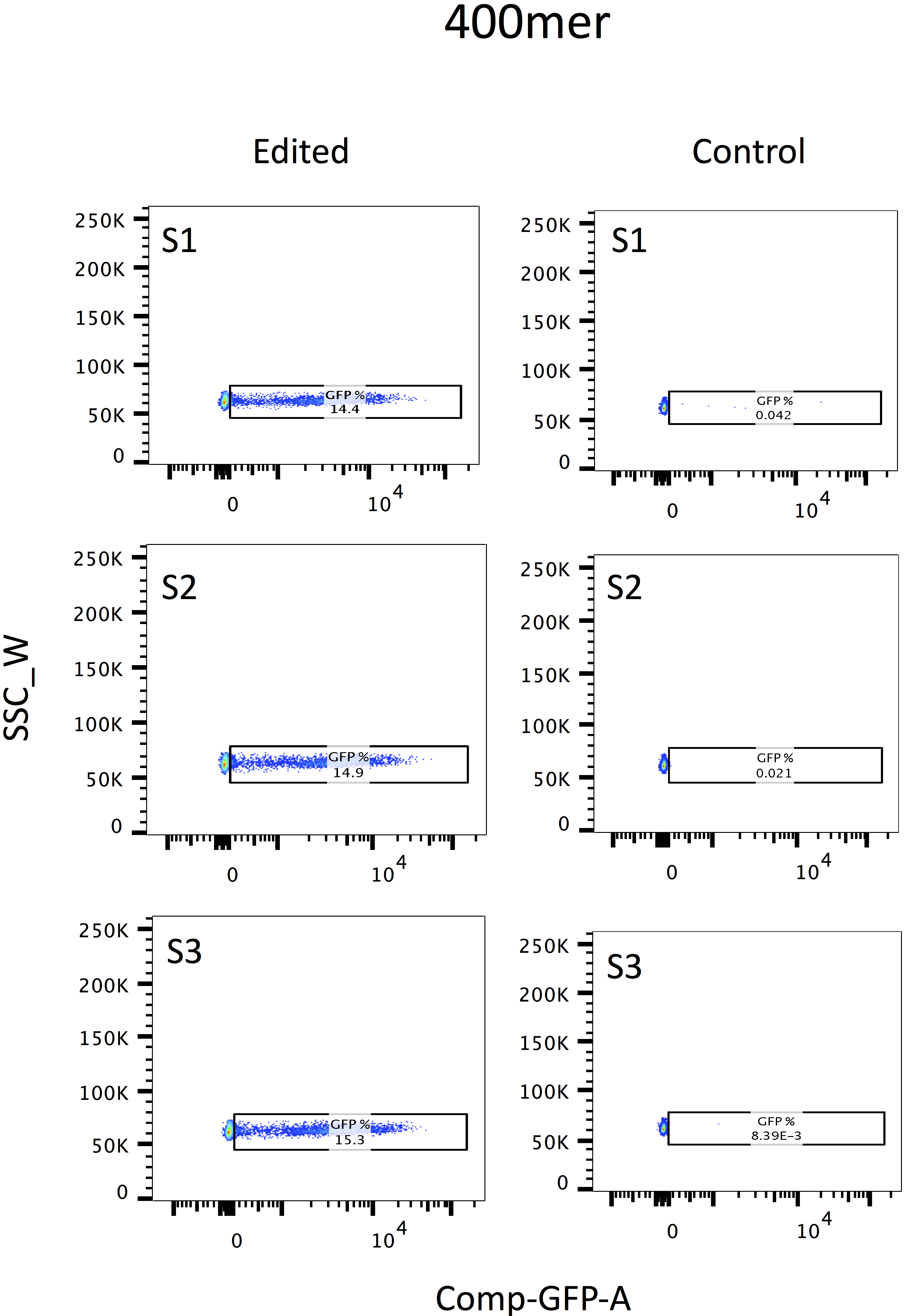

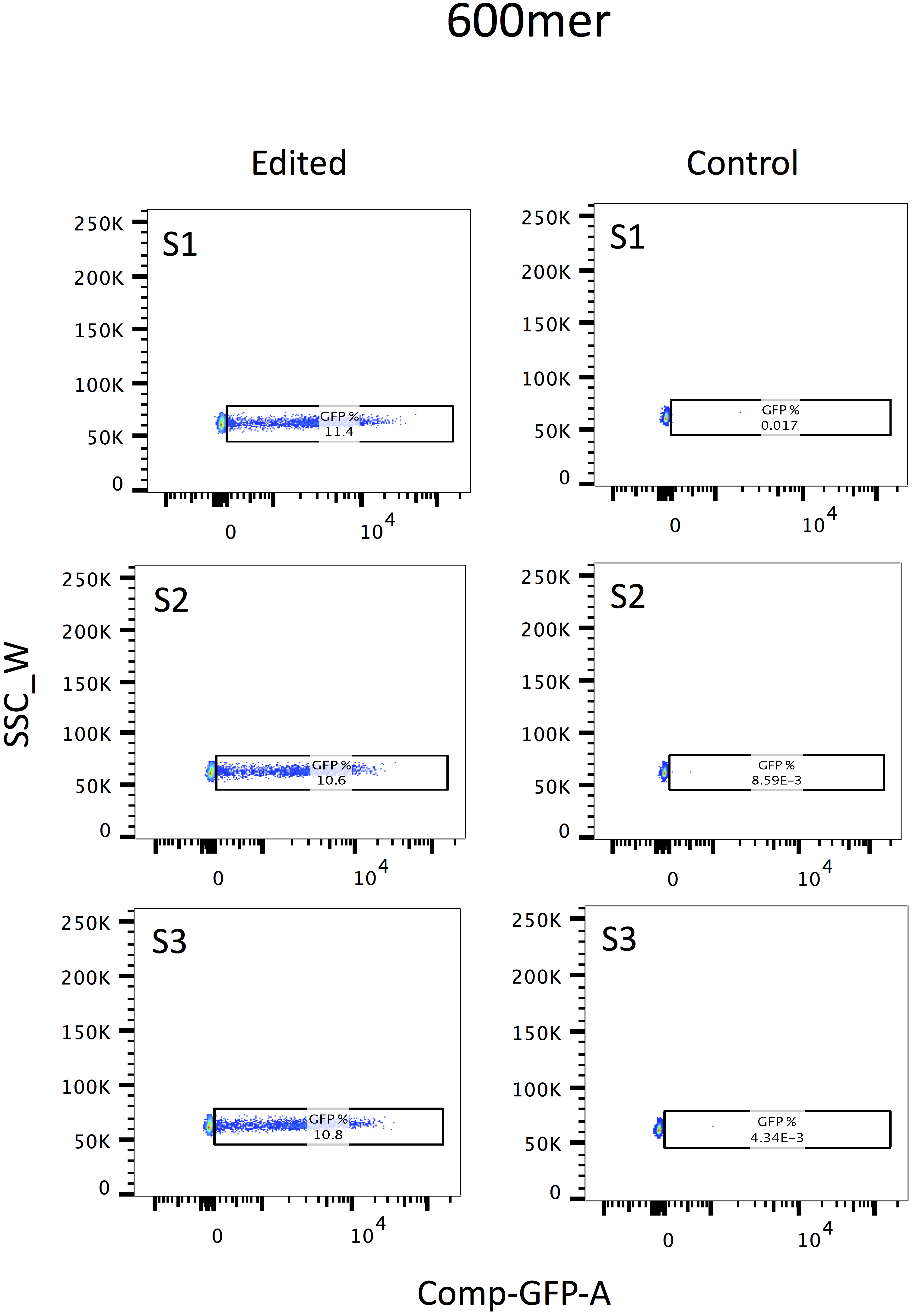

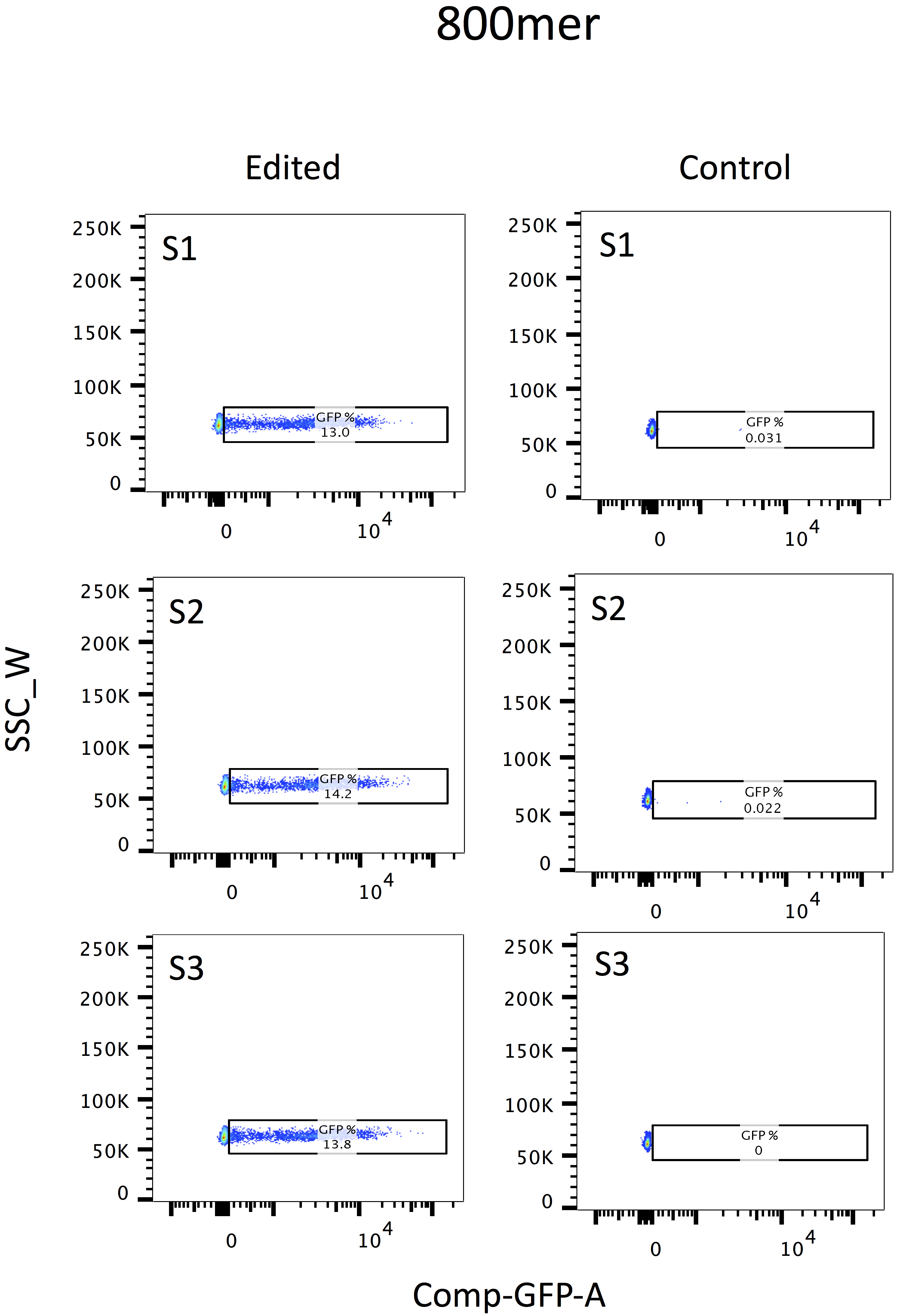

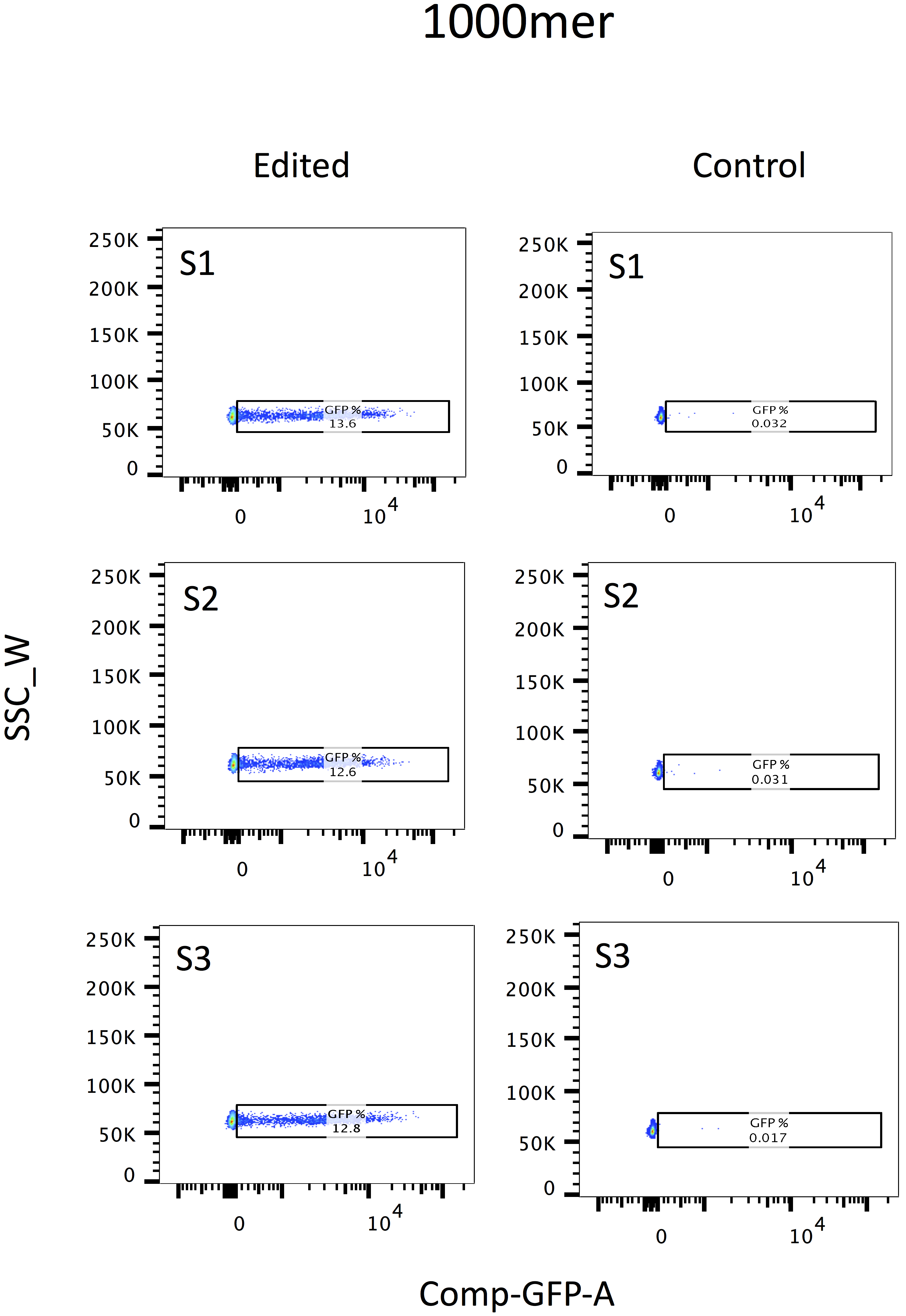
